# Genome based analyses reveals the presence of heterotypic synonyms and subspecies in Bacteria and Archaea

**DOI:** 10.1101/2020.12.13.418756

**Authors:** Munusamy Madhaiyan, Venkatakrishnan Sivaraj Saravanan, Wah-Seng See-Too

## Abstract

Term heterotypic synonym refers to different names have been associated with different type strains, however from the opinion of a bacteriologist, different names belongs to the same taxon and term subspecies refers to strains and genetically close organisms that were diverging phenotypically. In this study, sequenced and publicly available genomes in the Edgar 2.0 server were carefully analysed and based on high (>98 %) amino acid identity value, synonyms were putatively identified. The 16S rRNA gene sequence of those species were used for the construction of maximum likelihood based phylogenetic trees to infer the genetic closeness or distance by examining the tree topology and clustering of the organisms within clades. They were further subjected to overall genome related indices like digital DNA-DNA hybridization, average nucleotide identity to confirm the presence of synonyms or subspecies with phenotypic data support. The outcome of this polyphasic taxonomic re-analysis was identification of 40 later heterotypic synonyms and 13 subspecies spread over phylum *Actinobacteria*, *Bacteroidetes*, *Firmicutes*, *Nitrospirae*, *Proteobacteria* and *Thermotogae* and in domain *Archaea*.

## INTRODUCTION

A taxon refers to one or more elements, it can be of any taxonomic category from class to subspecies that are designated as nomenclatural type. The term type refers to the nomenclatural type strain designated as per the ICNP rule, it’s the element of the taxon with which the name is permanently associated as correct name or heterotypic synonym [1]. In simple terms, synonym refers to the same taxon under another scientific name, they usually come in pair or even swarms [2]. In old literature, they were named as objective synonyms and subjective synonyms which are referred in the recent times as homotypic and heterotypic synonyms [1]. In nutshell, homotypic synonym referring to more than one name associated with same type strain whereas heterotypic synonyms refer to different names have been associated with different type strains but based on the opinion of the bacteriologist concerned both names belong to the same taxon.

Heterotypic synonyms were first used in conjunction with junior synonyms, in a study conducted on *Helicobacter* type strains, where in *Helicobacter nemestrinae* [3] was identified as junior heterotypic synonym of *Helicobacter pylori* [4], further, the presence of later heterotypic synonyms were identified in different genera including *Janibacter*, *Weissella* and *Streptomyces* [5, 6, 7]. Conventionally, heterotypic synonyms were proposed based on the 16S rRNA gene analysis and DNA-DNA hybridization studies as in the case of *Janibacter* and *Weissella* [5, 6]. For instance, employing DNA-DNA hybridization technique on 13 *Streptomyces* species, two subspecies and 8 later heterotypic synonyms were identified [7]. A study provided polyphasic evidence for heterotypic synonym using fluorescent amplified fragment length polymorphism, DNA-DNA hybridization and API 50 CHL analysis on *Lactobacillus ferintoshensis* and *Lactobacillus parabuchneri* sharing similar genetic, biochemical and physiological features, this led to proposal of *L. ferintoshensis* as the later heterotypic synonym of *L. parabuchneri* [8]. Another study focused on using of multilocus sequence typing analysis to identify heterotypic synonyms, herein similarity of three housekeeping genes including *pheS*, *rpoA* and *atpA* were used to propose *Lactobacillus crypricasei* as the later heterotypic synonym of *Lactobacillus acidipiscis* [9]. In later studies, the heterotypic synonyms were identified when they attempted to characterize certain isolated strains or serendipitously discovered when attempted to validate a previously published strain [10, 11]. A new dimension in the study of heterotypic synonyms was the suggestion of the use of average nucleotide identity (ANI), since the next generation sequencing provided a rapid and cost effective way of obtaining the whole genome sequences, a proposal for integrating the genomics as a polyphasic component of the Bacteria and Archaea taxonomy and systematics was mooted out [12], ANI values and comparative genomic analysis had become an inevitable part in the proposal of heterotypic synonyms in *Neisseria*, *Deinococcus* and *Bacillus* [13, 14, 15].

The term subspecies denote the strains and genetically similar organism that are phenotypically divergent [16, 17]. Previously, delineation of subspecies was based on qualitative measurement of phenotypic characters rather than examining evolutionary distance or 16S rRNA gene similarity. However, the current approach in the taxonomic assignment of subspecies was based on the overall genome related indices such as digital DNA-DNA hybridization (dDDH) of 70-79 % [18].

The advent of genome sequencing technologies was reflected in bacterial systematics mainly in reclassification of the taxa at different hierarchy level, such kind of proposals were attempted in phylum *Actinobacteria*, *Bacteroidetes*, and *Alphaproteobacteria* [19, 20, 21]. As an exemplar, within the taxa of *Actinobacteria*, by calculating intergenomic dDDH values, 29 new later heterotypic synonyms, 8 new subspecies were proposed [19], in the phylum *Bacteroidetes* 6 later heterotypic synonyms and 5 new subspecies were proposed [20]. The recent analysis of the phylum *Alphaproteobacteria* revealed the presence of 33 later heterotypic synonyms, 12 new subspecies and 2 new species [21]. The present study focusses on whole genome-based analyses to identify heterotypic synonyms in the sequenced and publicly available bacterial genomes. The idea behind this work originated when species of certain publicly available genomes in the Edgar 2.0 server [22] showed high amino acid similarity when their genomes were compared. Previous studies show that genome analysis clearly envisage that strains from same species share an amino acid identity (AAI) of > 95 % [23]. Based on this notion, on total 12479 genomes comprising of 548 genera housed within 227 families were examined. Different species showing > 98 % AAI similarity were assumed to contain heterotypic synonym or subspecies and they were selected and used in the present study. Alternatively, genome results of certain recently published type strains when critically examined, the possibility of heterotypic synonymy was predicted. These putatively identified strains were also subjected to phylogenetic analysis, overall genome related indices (OGRI) such as dDDH and average nucleotide identity (ANI) analyses and their phenotypic traits were also considered to propose a holistic taxonomic framework for the heterotypic synonyms and subspecies present in different genera.

## METHODS

In this study, 95 type strains of different species of Bacteria and Archaea whose genome shared > 98 % AAI were selected from the Edgar 2.0 server and certain recently published genome contiaing putative heterotypic synonmy were downloaded from NCBI. Strains of these selected genomes were distributed among phylum *Actinobacteria, Bacteroidetes, Firmicutes, Nitrospirae, Proteobacteria, Thermotogae* and Domain *Archaea*, their genome details are presented in Table S1.

For the selected strains, the pairwise 16S rRNA gene sequence similarity was worked out in a phylum wise manner using the EzBioCloud database [24]. The Clustal W tool of MEGA 7 was used for alignment of the sequences in an individual phylum level [25], distance between the sequences was calculated using the Kimura correction in a pairwise deletion manner [26] and maximum likelihood-based treeing program was used to reconstruct the type strains phylogeny using the MEGA 7 software with 1000 non-parametric bootstrap replicates.

For OGRI analysis, 95 genome sequences of different type species were retrieved from EzBioCloud database /GenBank. Using the retrieved genomes, OGRI analysis including Genome distance in the form of digital DNA-DNA hybridization (dDDH) values calculated by the Genome-to-Genome Distance Calculator (GGDC) version 2.1 (http://ggdc.dsmz.de) with BLAST+ and with the recommended formula [27]. OrthoANI, that measures similarity between two genome sequences was calculated using Oat 0.93.1 [27] with USEARCH (https://www.ezbiocloud.net/tools/orthoaniu). Average amino acid identities (AAI) was calculated using the EDGAR server 2.3 (https://edgar.computational.bio.uni-giessen.de). For certain pairs retrived from database, the pairwise average AAI was calculated using the AAI workflow with default settings in CompareM v0.0.23 (https://github.com/dparks1134/CompareM).

## RESULTS AND DISCUSSION

When the genomes of the phyla were analysed, presence of synonyms or subspecies were putatively identified by 16S rRNA gene sequence based phylogenetic tree topology, the gene similarity values and clustering of the organismsm within the clades. The selected species pairs or taxa were further subjected to dDDH [27], to underpin the subspecies or synonynm with a threshold values of 70 - 79 % or > 80 % respectively [17, 18]. The complete list of subspecies and heterotypic synonyms identified employing these criteria are listed in the Table 1.

**Table 1.**
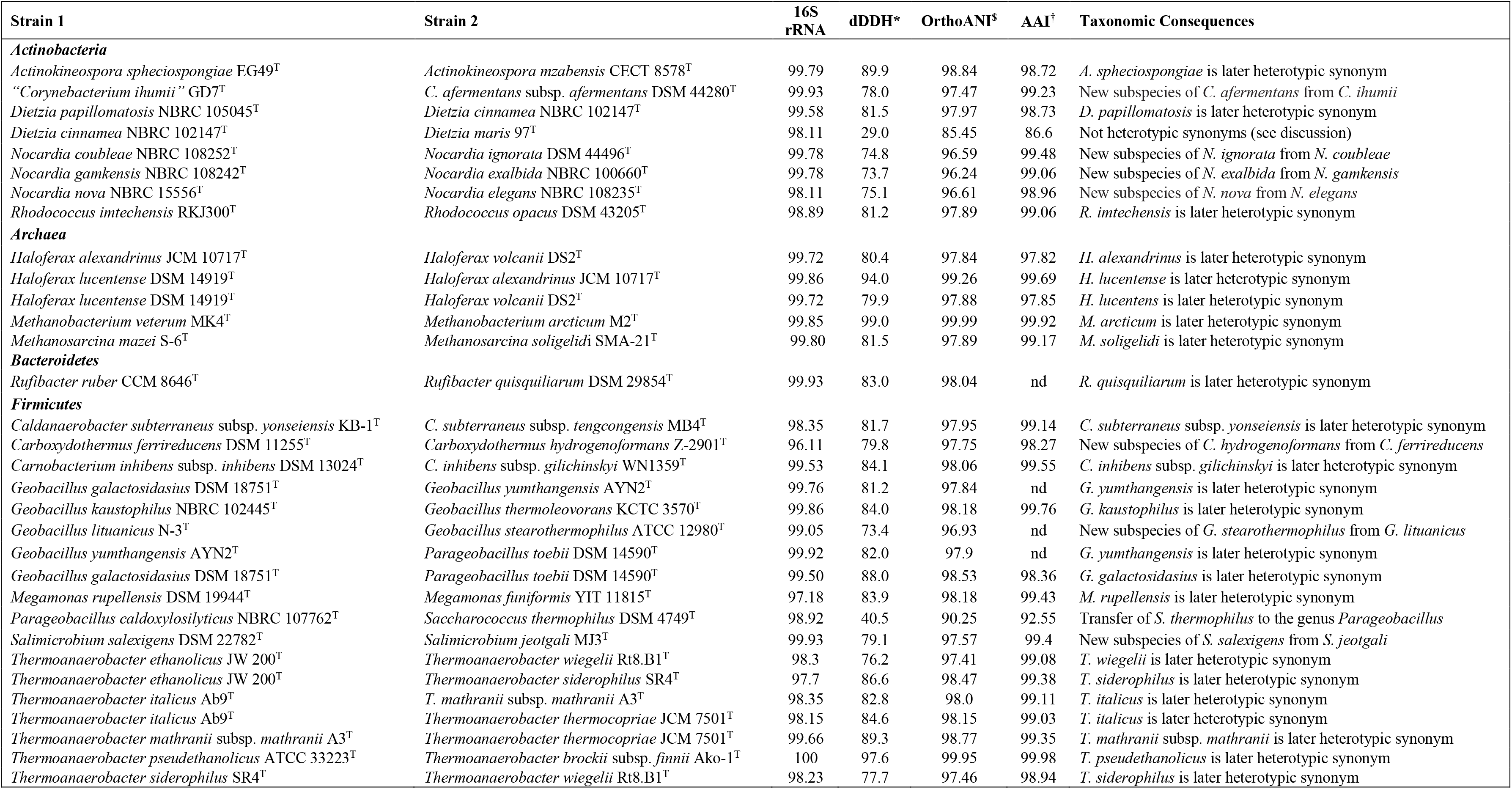
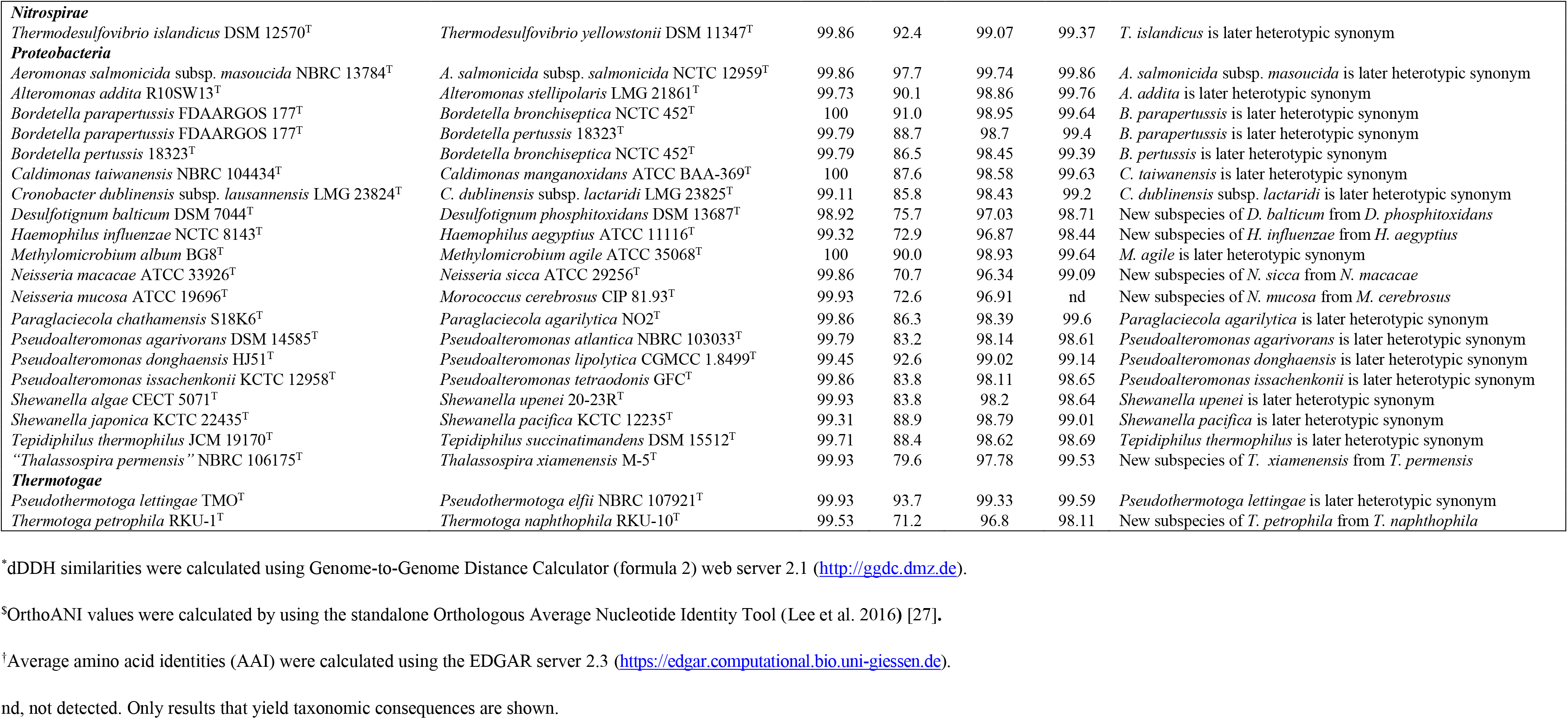
16S rRNA gene sequence percent identity and overall genome-relatedness indices for specific clades in the phylum *Actinobacteria, Bacteroidetes, Firmicutes, Nitrospirae, Proteobacteria, Thermotogae* and in domain *Archaea*.

### Actinobacteria

*Actinobacteria* is one important phylum that was analysed for the presence of heterotypic synonyms, the putative synonymous strains in different species were selected based on the high AAI (>98 %) values of their genomes. The genetic closeness among the different species pairs indicating the presence of heterotypic synonyms or subspecies was preliminarily identified based on high similarity per centage of their 16S rRNA gene sequences (Table 1).

For instance, a genetically close pair consists of *Actinokineospora spheciospongiae* and *Actinokineospora mzabensis* showing high similarity and bootstrap support in the 16S rRNA gene analysis (Fig. 1). When it was subjected to the OGRI metrics (Table 1), this pair shared dDDH of 89.9 %, their genome similarity was also reflected in the biochemical properties, as only a few differences in mycelial pigments and carbon source utilization was observed between the strains (Table S2). These data mooted the proposal of *Actinokineospora spheciospongiae* as the later heterotypic synonym of *Actinokineospora mzabensis*. Further, when a *Corynebacterium* clade was analysed, *Corynebacterium ihumii* recorded high dDDH value of 76.1 and 78 % with *Corynebacterium afermentans* subsp. *lipophilum* HSID17239 and *Corynebacterium afermentans* subsp. *afermentans* DSM 44280^T^ respectively, phenotypically most of the traits are common between these strains (Table S3). The dDDH value supported the reduction of taxon *Corynebacterium ihumii* as *C. afermentans* subsp. *ihumii*. However, *Corynebacterium ihumii* was not validly published and the subspecies was not proposed as per the taxonomic rules and guidelines [29]. This clade also comprised of another pair of *Corynebacterium pilbarense* and *C. ureicelerivorans* (Fig. 1) and its dDDH value was less than 70 % retaining their species status. A recent study by Nouioui et al. [19] proposed *Dietzia cinnamea* as the later heterotypic synonym of *Dietzia maris*, when the dDDH value was re-analysed it shared only 29 %, erroneous results obtained may be attributed to the incorrect strain of *D. maris* DSM 43672 used in that study. Thus *D. cinnamea* retaining its species status, but, when the genome of *D. cinnamea* was compared with *D. papillomatosis*, dDDH was 81.55 %, phenotypically also both the strains were common in morphological feaatures including their colony colour to substrate utilization profile (Table S4), but *D. maris* strain was phenotypically different from *D. cinnamea* and *D. papillomatosis*. Thus, the OGRI combined with phenotypic features supported the proposal of *Dietzia papillomatosis* as the later heterotypic synonym of *Dietzia cinnamea*. A monophyletic clade of *Nocardia* containing certain species sharing high 16S rRNA gene similarity (Table 1 & Fig. 1) was noticed, all those three pairs were analysed for the dDDH to ascertain their taxonomic position (Table 1), all strains recorded within a range of 73.7 - 75.2 %, which coincides with the value (70-79 %) proposed as cutoff to delineate subspecies [18]. Further presence of similar phenotypic properties is noticed within those pairs. For instance, the pair consist of *N. exalbida* and *N. gamkensis* shared similarities in the degradation of substrates including arbutin, esculin adenine, casein, hypoxanthine and xanthine (Table S5). The other pair comprising of *Nocardia ignorata* and *Nocardia coubleae* when analysed, similarities in abiotic factors (temperature, pH and NaCl concentration) affecting the growth was observed (Table S6). The third pair housing *N. elegans* and *N. nova* were biochemically similar in utilization of carbon sources as sole carbon source and energy (Table S7). The dDDH value combined with the phenotypic traits supported the proposal of three different subspecies within *Nocardia* clade namely *Nocardia exalbida* subsp. *gamkensis* subsp. nov.*, Nocardia ignorata* subsp. *coubleae* subsp. nov. and *Nocardia nova* subsp. *elegans* subsp. nov. In the *Rhodococcus* clade, genetic closeness of a pair was identified through 16S rRNA based phylogenetic tree (Fig. 1), when it was subjected to dDDH analysis, values supporting the claim of a heterotypic synonym was observed (Table 1). In fact, phenotypically, most of the biochemical features are common between these strains (Table S8). This led to the proposal of *Rhodococcus imtechensis* is later heterotypic synonym of *Rhodococcus opacus*.

**Fig. 1.**
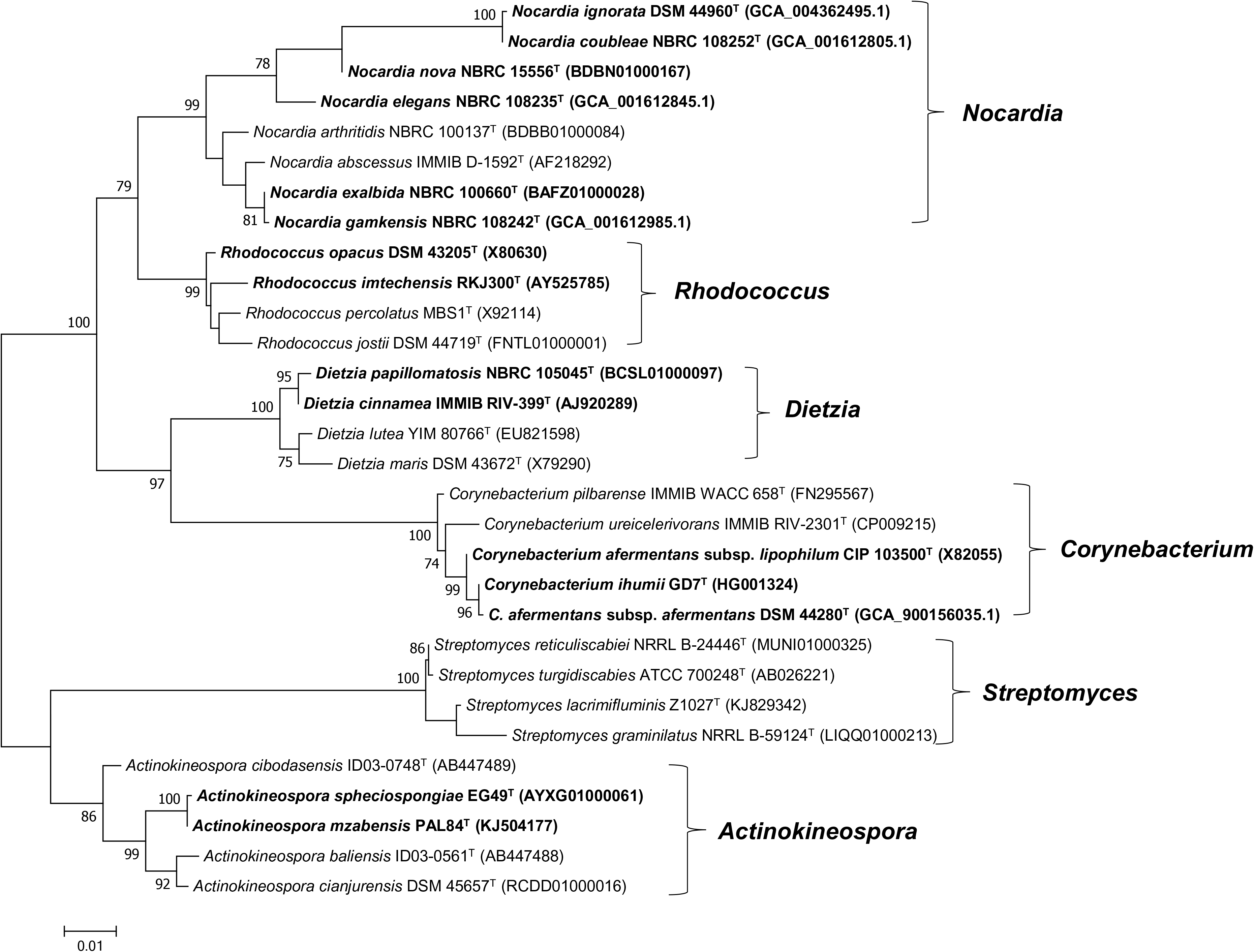
Phylogenetic analysis based on the 16S rRNA gene sequences comparison highlighting the position of 8 heterotypic synonyms from 4 different genera and other closely related strains belongs to the phylum *Actinobacteria*. The numbers at the nodes are given as percent and represent the levels of bootstrap support (>70%) based on maximum likelihood analyses of 1000 re-sampled data sets. GenBank accession numbers are indicated in parentheses. Heterotypic synonyms of the type species are marked in bold. Sequences of the strains were retrieved from EzBioCloud. Scale bar, 0.01 nt substitutions per position.

### Archaea

Presence of heterotypic synonyms were putatively selected in Domain *Archaea* through preliminary examination of AAI value in genome belong to different species. Those showing high AAI (>98 %) were taken up for further OGRI analysis. Presence of heterotypic synonym was recognized within the clade of *Haloferax* (Fig. 2), high 16S rRNA gene similarity (Table 1) was observed between three strains including *Haloferax lucentense*, *Haloferax alexandrinusand Haloferax volcanii*, dDDH analysis revealed the genome similarity of those strains (Table 1). Phenotypically, all the strains are oxidase positive and shared a similar concentration of NaCl requirement (1-4.5 % w/v) pH and temperature optimum for growth. All strains produced H_2_S when grown on thiosulfate medium and formed acid from arabinose and none of the strains hydrolysed starch and casein, but variable results observed for gelatin hydrolysis (Table S9). Based on the 16S rRNA gene sequence similarity, dDDH value and biochemical features, *Haloferax lucentense* and *Haloferax alexandrinus* are proposed as the heterotypic synonyms of *Haloferax volcanii*.

**Fig. 2.**
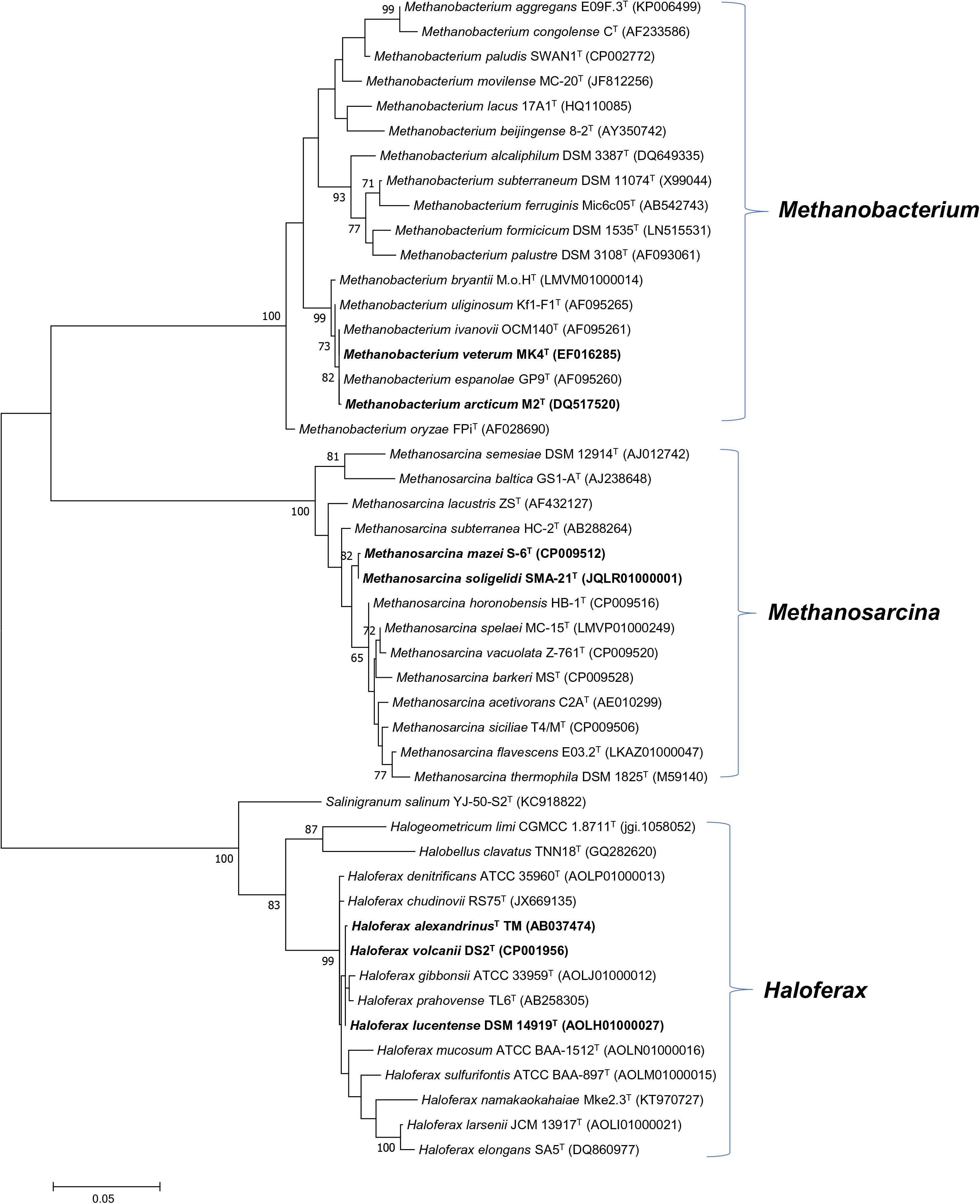
Maximum- likelihood phylogenetic tree based on the 16S rRNA gene sequences of taxa belonging to domain *Archaea*, the tree depicts the phylogenetic position of 5 heterotypic synonyms identified in 3 different genera and their closely related members. Bootstrap values are expressed as percentages of 1000 replications and are shown at branch nodes (>70 %). GenBank accession numbers are indicated in parentheses. Heterotypic synonyms of the type species are marked in bold. Sequences of the strains were retrieved from EzBioCloud. Scale bar, 0.05 nt substitutions per position.

Another clade comprising of *Methanobacterium* contained two species that may be genetically close based on 16S rRNA gene similairty (Fig. 2). Those species were subjected to dDDH analysis (Table 1), high genetic similarity was noticed between *Methanobacterium veterum* and *Methanobacterium arcticum*. Critical examination of their phenotypic traits revealed the fact that both the strains share similar temperature, pH and NaCl concentration ranges supporting growth (Table S10). The 16S rRNA gene sequence-based analysis (Fig. 2) revealed that other *Methanobacterium* species present within the clade may also genetically similar, but currently genome data was lacking for all those species except *Methanobacterium bryantii*, which showed only 48 % dDDH value with *M. veterum* and *M. arcticum*. Considering together the OGRI and biochemical properties, *Methanobacterium arcticum* is proposed as a later heterotypic synonym of *Methanobacterium veterum*. Similarly, the heterotypic synonyms were identified in *Methanosarcina* based on high 16S rRNA gene similarity between *Methanosarcina soligelidi* and *Methanosarcina mazei*, it was also reflected in their genome based dDDH analysis (Fig. 2 & Table 1), the observed results were supported by phenotypic properties including abiotic factors (temperature, pH and NaCl concentration) and substrate utilization pattern, interestingly, both shared similar archaeal lipids in their membrane (Table S11). These polyphasic data supported the proposal of *Methanosarcina soligelidi* as the later heterotypic synonym of *Methanosarcina mazei*.

### Bacteroidetes

In *Bacteroidetes*, species pair consists of *Rufibacter ruber* and *Rufibacter quisquiliarum* were similar in their genome which was evident from the values recorded in dDDH (Fig. 3 & Table 1). Interestingly, in between these strains, abiotic factors such as temperature, pH and NaCl concentrations favouring growth are common with few differences in carbon source assimilation (Table S12). These polyphasic evidences support the proposal of *R. quisquiliarum* as the later heterotypic synonym of *R. ruber*.

**Fig. 3.**
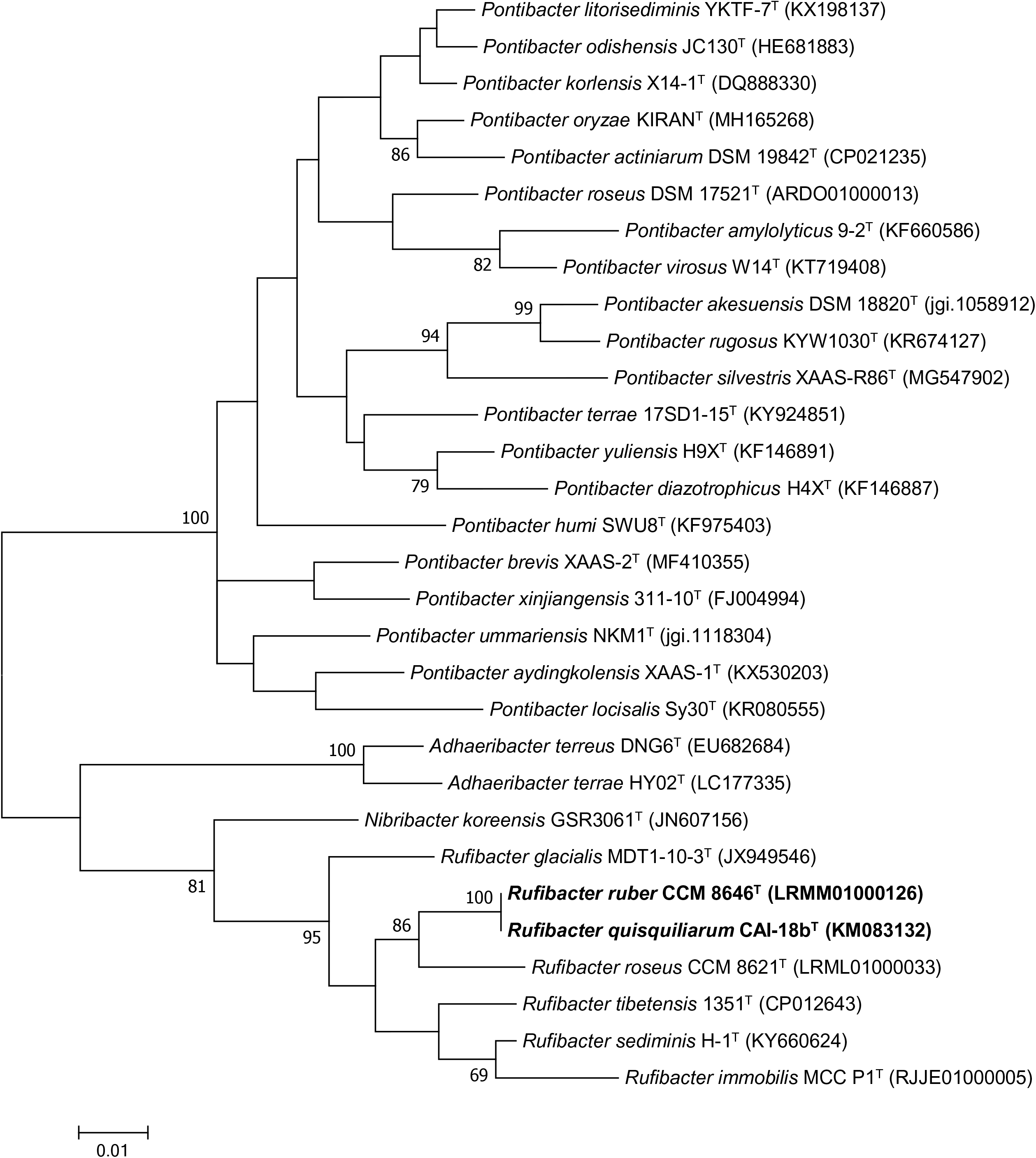
*Bacteroidetes* members 16S rRNA gene phylogeny reconstructed using Maximum- likelihood method, it shows the heterotypic synonomous strains *Rufibacter ruber* CCM 8646^T^ and *Rufibacter quisquiliarum* CAI-18b^T^ and its top 30 closest relatives with validly published name. The numbers at the nodes are given as percent and represent the levels of bootstrap support (>70%) based on maximum likelihood analyses of 1000 re-sampled data sets. GenBank accession numbers are indicated in parentheses. Scale bar, 0.01 nt substitutions per position.

### Firmicutes

A subspecies pair consists of *Caldanaerobacter subterraneous* subsp. *yonseiensis* and *Caldanaerobacter subterraneous* subsp. *tengcongensis* showed high 16S rRNA gene similarity (Fig. 4), they also shared dDDH and ortho ANI values of 81.7 and 98.0 % respectively. The genome level similarity was also reflected in phenotypic properties with very few variable traits (Table S13). Another organism that was phylogenetically close and found within this clade was *Thermoanaerobacter keratinophilus*, which may be a taxon belonging to *Caldanaerobacter*, but currently it was not validly published, due to the fact that only a single culture collection (*T. keratinophilus* DSM14007^T^) deposition proof was available and the non-availability of the genome details also hampered its re-classification. Based on the OGRI metrics analysed *Caldanaerobacter subterraneus* subsp. *yonseiensis* was proposed as the heterotypic synonym of *C. subterraneus* subsp *tengcongensis*.

**Fig. 4.**
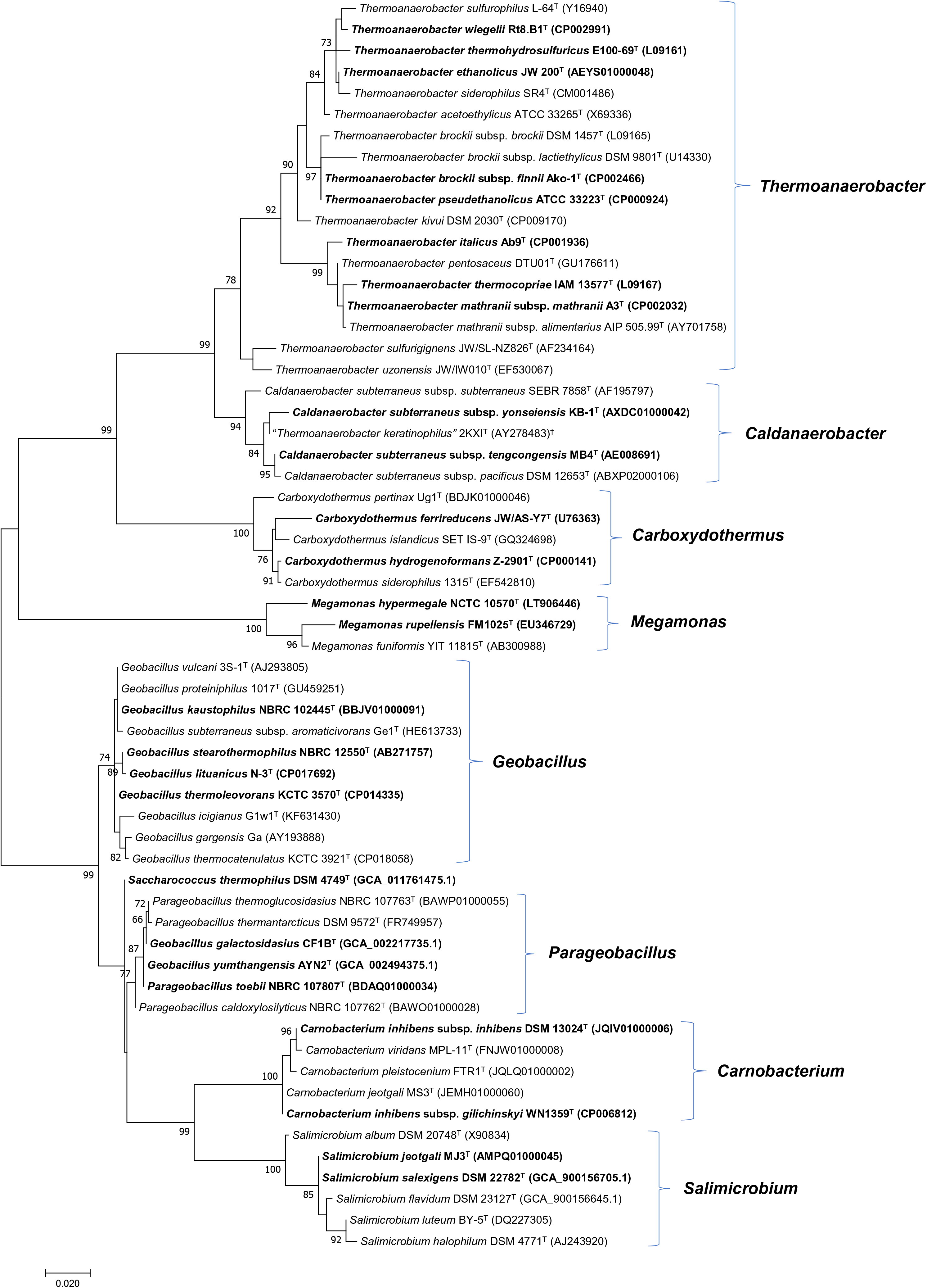
Maximum-likelihood tree based on 16S rRNA gene sequences of 18 heterotypic synonyms belongs to 8 different genera within the phylum *Firmicutes.* The numbers at the nodes are given as percent and represent the levels of bootstrap support (>70%) based on maximum likelihood analyses of 1000 re-sampled data sets. GenBank accession numbers are indicated in parentheses. Heterotypic synonyms of the type species are marked in bold. Scale bar, 0.01 nt substitutions per position.

Another pair consists of *Carboxydothermus hydrogenoformans* and *Carboxydothermus ferrireducens* sharing 96.1 % 16S rRNA gene similarity (Fig. 4) and shared 79.8 and 97.8 % of dDDH and ortho ANI respectively (Table 1). Biochemically there is not much difference existing between these two species, except the utilization of ferric citrate as electron acceptor during chemolithotrophic growth of cells with CO as electron donor (Table S14). The 16S rRNA analysis also shows the presence of *Carboxydothermus siderophilus* within this clade (Fig. 4), its taxonomic position was not analysed due to the non-availability of the genome data. Thus, the dDDH recorded within the range of 70-79 % supported the reduction of taxon *Carboxydothermus ferrireducens* to *Carboxydothermans hydrogenoformans* subsp. *ferrireducens* subsp. nov.

The next pair analysed within the *Firmicutes* consists of *Carnobacterium inhibens* subsp. *inhibens* and *Carnobacterium inhibens* subsp. *gilichinskyi* sharing 16S rRNA gene similarity of 99.4 % and their dDDH and ortho ANI recording 84.1 and 98.1 % respectively (Table 1). Both strains also required a wider temperature (0-37 ^ο^C) and pH (5.5-9.0) for growth, with a little variability noticed in the utilization of substrates (Table S15). On careful observation of the 16S rRNA gene based phylogenetic tree (Fig. 4), this clade has also nested *Carnobacterium viridans*, *C. pleistocenium* and *C. jeotgali*. Especially *C. pleistocenium*, *Carnobacterium inhibens* subsp. *gilichinskyi* and *C. jeotgali* shared a uniform bootstrap support, that lead to analysis of their genomes to identify heterotypic synonyms or subspecies. However, all the three strains have recorded <70 % dDDH retaining their species status. The OGRI including dDDH and ortho ANI strongly supported proposing *Carnobacterium inhibens* subsp. *inhibens* as the later heterotypic synonym of *Carnobacterium inhibens* subsp. *gilichinskyi*.

Another important genus analysed for the presence of heterotypic synonyms or subspecies in *Firmicutes* was *Geobacillus*. On analyses of the twelve species of *Geobacillus* and four *Parageobacillus* for OGRI metrics of dDDH and Ortho ANI. The two *Geobacillus* pairs consist of *G. kaustophilus*; *G. thermoleovorans* and *G. lituanicus*; *G. stearothermophilus* shared a high level of 16S rRNA gene similarity (Fig. 4), and the dDDH and ortho ANI ranged between (73.4-88.0 %) and (96.9-98.5 %) respectively (Table 1). The careful examination of the phenotypic traits showed only little variation within the taxa (Table S16) strongly supporting the claim that *Geobacillus kaustophilus* is the later heterotypic synonym of *Geobacillus thermoleovorans*. Another pair that consist of *Geobacillus lituanicus* and *Geobacillus stearothermophilus* has recorded dDDH value of 73.4 %, Interestingly, both the species shared a common temperature regime (Table S17), with optimum temperature requirement of 50 ^ο^C for growth [30, 31, 32]. The dDDH recorded and similarity in phenotypic traits supported the subspecies delineation [18], and proposal of different subspecies namely *Geobacillus stearothermophilus* subsp. *stearothermophilus* subsp. nov. While these pairs were analysed, a pair of *Geobacillus* comprising of *Geobacillus yumthangensis* and *Geobacillus galactosidasius* were found to be clustered within the *Parageobacillus* clade (Fig. 4) in fact both the strains showed high genomic similarity indices (Table 1) in the form of dDDH (82- 88 %) and ANI (97.9 - 98.5 %) with *Parageobacillus toebii*, variability in the biochemical traits was observed between the three strains are: production of acid from cellobiose, galactose, ribose, glycerol, lactose and rhamnose, variation was recognized within the carbon source utilization also. But, the abiotic factors (temperature, pH and NaCl concentration) required for these strains are similar (Table S18). Thus the, 16S rRNA based topology with genome indices and biochemical data strongly supported the proposal of *Geobacillus yumthangensis* and *Geobacillus galactosidasius* as later heterotypic synonyms of *Parageobacillus toebii*, in fact *Parageobacillus toebii* was a homotypic synonym of *Geobacillus toebii* [33].

*Megamonas* and *Salimicrobium* are the other two genera of *Firmicutes* that were analysed for the OGRI metrics. Genome similarity of their strains can be understandable by the critical analysis of the dDDH and ANI of the *Megamonas* pair (*M. rupellensis* and *M. funiformis*) and *Salimicrobium* pair (*S. salexigens* and *S. jeotgali*) (Table 1). One more taxon (*M. funiformis*) was associated with the *Megamonas* clade (Fig. 4) but, currently its genome sequence was unavailable. Phenotypic similarity within these two pairs are also high, only a little difference in the form of carbon substrate utilization and acid production from carbon sources are observed (Table S19 - S20). Thus, the OGRI analyses supported the proposal of *Megamonas rupellensis* as the later heterotypic synonym of *Megamonas funiformis*, previously also a synonym (*M. hypermegas*) was identified in this genus [34]. In the *Salimicrobium* clade analysed, *S. jeotgali* was reduced in taxon and proposed as *Salimicrobium salexigens* subsp. *jeotgali* subsp. nov. a novel subspecies as per the previously suggested threshold value [18].

A paraphyletic clade comprised of eight *Thermoanaerobacter* species (Fig. 4) was subjected to OGRI analyses (Table 1), subsequently, their biochemical characteristics was also compared, a few differences in the form of motility and carbon substrate utilization was observed (Table S21) The OGRI and phenotypic evidences supported the proposal of *Thermoanaerobacter italicus* and *Thermoanaerobacter mathranii* subsp. *mathranii* as the later heterotypic synonyms of *Thermoanaerobacter thermocopriae*. *Thermoanaerobacter pseudethanolicus* as the later heterotypic synonym of *Thermoanaerobacter brockii* subsp. *finnii*. Finally, *Thermoanaerobacter wiegelii* and *Thermoanaerobacter siderophilus* together constitute the later heterotypic synonyms of *Thermoanaerobacter ethanolicus*.

### Nitrospira

The phylum *Nitrospira* comprised of three valid genera namely *Leptospirillum*, *Nitrospira* and *Thermodesulfovibrio* with *Thermodesulfovibrio* as the only genus containing five species. A species pair of *Thermodesulfovibrio* was examined owing to high AAI value and the analysis of its dDDH revealed the genome similarity between these two species (Table 1 & Fig. 5). Phenotypically, these strains utilized pyruvate, hydrogen (plus acetate) and formate (plus acetate) as both electron donor and carbon source in the presence of sulfate as terminal electron acceptor and in addition they also utilized sulfate and thiosulfate as electron acceptors. However, variability was noticed in utilization of sulfite and nitrate as terminal electron acceptors (Table 22). Ecologically, both the strains were recovered from hot spring habitats [35, 36]. These polyphasic evidences led to proposal of *Thermodesulfovibrio islandicus* as the later heterotypic synonym of *Thermodesulfovibrio yellowstonii*.

**Fig. 5.**
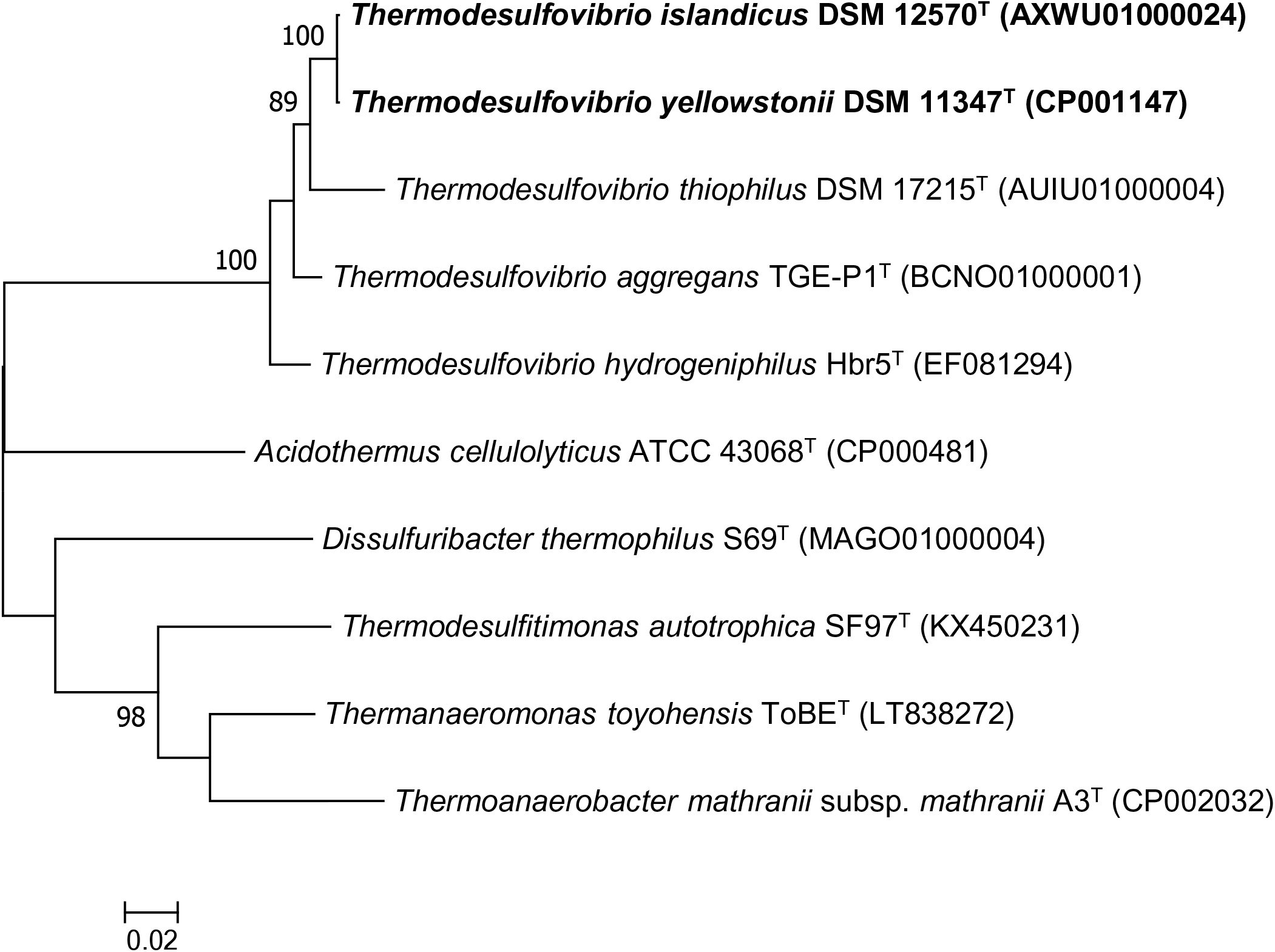
Maximum likelihood tree based on 16S rRNA gene sequences showing the phylogenetic relationship of strains *Thermodesulfovibrio yellowstonii* DSM 11347^T^ and *Thermodesulfovibrio islandicus* DSM 12570^T^ and other closely related strains belonged to the phylum *Nitrospirae*. The numbers at the nodes are given as percent and represent the levels of bootstrap support (>70%) based on maximum likelihood analyses of 1000 re-sampled data sets. GenBank accession numbers are indicated in parentheses. Heterotypic synonyms of the type species are marked in bold. Scale bar, 0.02 nt substitutions per position.

### Proteobacteria

*Proteobacteria* commonly refers to Gram negative bacteria, teems with versatile metabolic capability and omnipresent in nature. The genetic closeness between several species those belong to *Proteobacteria* were depicted clearly in the 16S rRNA gene sequence based phylogenetic tree (Fig. 6).

All the *Aeromonas* taxa analysed in this study clustered together as single clade (Fig. 6) with high (100 %) bootstrap support, this prompted to check the dDDH values for all the species within that clade, however, genome data was available only for *Aeromonas salmonicida* subsp. *pectinolytica* that recorded dDDH of < 70 %, retaining its subspecies status. Interestingly, based on the 16S rRNA gene sequence similarity, a *Haemophilus piscium* was found within the *Aeromonas* clade, and it was proposed as *Aeromonas salmonicida* subsp. *piscium* comb. nov. A subspecies pair consist of *Aeromonas salmonicida* subsp. *masoucida* and *Aeromonas salmonicida* subsp. *salmonicida* showed high AAI, so they were subjected for dDDH analysis (Table 1), phenotypically these strains sharing similar phenotypic and biochemical traits (Table S23). *A. salmonicida* subsp. *masoucida* was proposed as a later heterotypic synonym of *Aeromonas salmonicida* subsp. *salmonicida*. Currently, *Aeromonas salmonicida* comprised of five subspecies and the proposed one is the first heterotypic synonym of this genus.

**Fig. 6.**
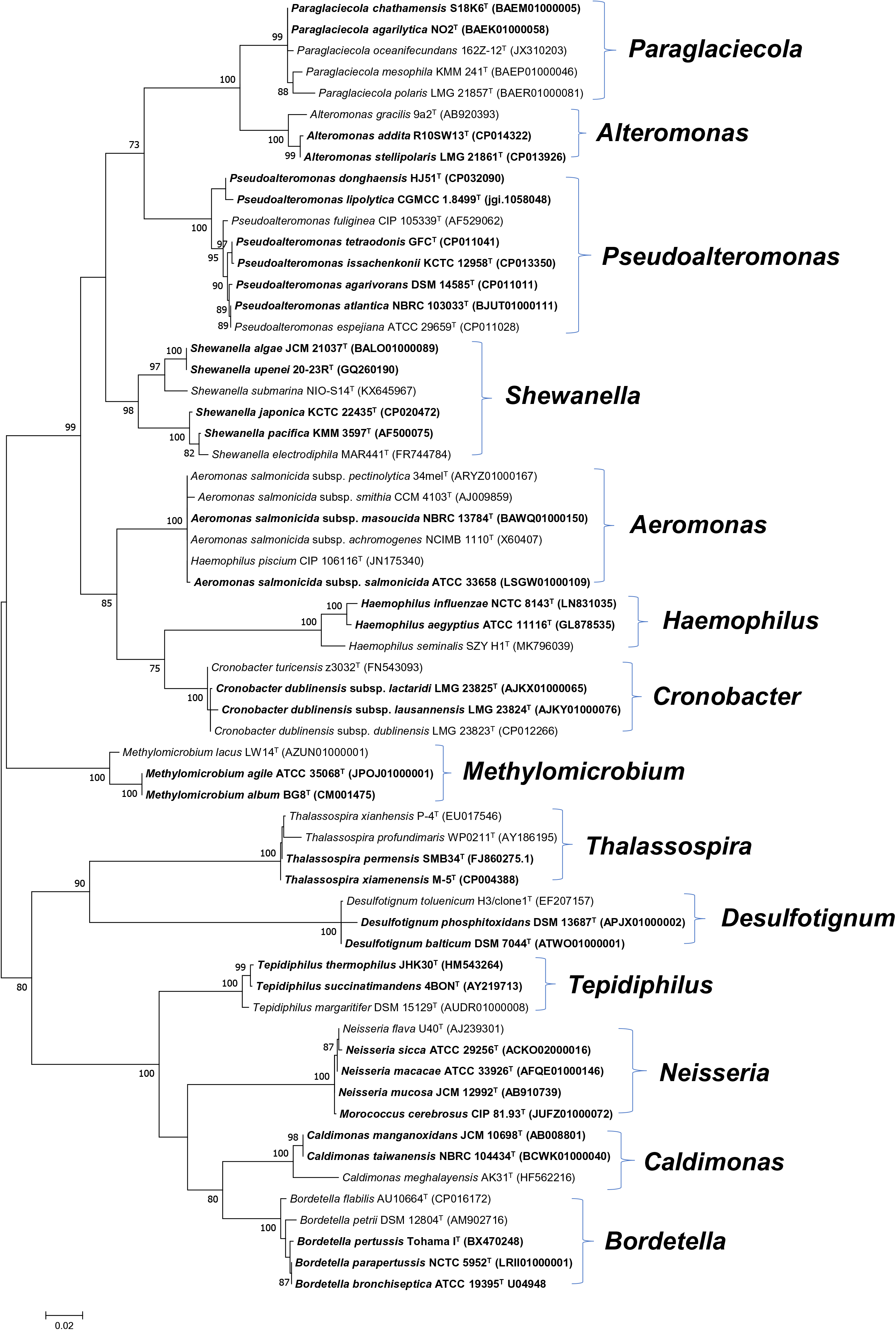
Maximum-likelihood tree based on 16S rRNA gene sequences of 27 heterotypic synonyms that belongs to 9 different genera within the phylum *Proteobacteria.* The numbers at the nodes are given as percent and represent the levels of bootstrap support (>70%) based on maximum likelihood analyses of 1000 re-sampled data sets. GenBank accession numbers are indicated in parentheses. Heterotypic synonyms of the type species are marked in bold. Scale bar, 0.02 nt substitutions per position.

A pair of *Alteromonas* species (*A. addita* and *A. stellipolaris*) was subjected for the dDDH analysis (Table 1). Both these strains shared a similar abiotic factor (temperature and NaCl) requirements for their growth and metabolism (Table S24). Ecologically, both strains share similar environment as they were isolated from sea water samples [37, 38]. The genome metrics and phenotypic evidences supported the proposal of *Alteromonas addita* as the later heterotypic synonym of *Alteromonas stellipolaris*. A prevous phylogenetic analysis study of the *Alteromonas* taxa, lead to transfer of 11 species to the newly proposed genus *Pseudoalteromonas*, leading to large numbers of synonyms in this genus [39].

One of the important genus from the perspectives of clinical microbiology was *Bordetella* [40], whose taxa were screened for heterotypic synonym owing to high 16S rRNA gene similarity recorded within three species of *Bordetella* (*Bordetella parapertussis, Bordetella bronchiseptica* and *Bordetella pertussis*) (Table 1), their genome were analysed for heterotypic synonyms or subspecies (Fig. 6). Deduction of high dDDH values (Table 1) between these strains suggested that at least two species among them may be heterotypic synonyms, further the analyses of phenotypic properties also supported this claim as certain carbon utilization and enzyme activity patterns are similar among all these species (Table S25). Genome based OGRI metrics combined with biochemical features supported the proposal of *Bordetella parapertussis* and *Bordetella pertussis* as the later heterotypic synonyms of *Bordetella bronchiseptica*.

Further, 16S rRNA gene similarity-based closeness was noticed in the following genera namely *Caldimonas, Cronobacter, Desulfotignum, Haemophilus, Methylomicrobium, Paraglaciecola, Tepidiphilus* and *Thalassospira* (Fig. 6). Their species pairs were analysed for OGRI metrics like dDDH and ANI to identify heterotypic synonym or subspecies within their species clades (Table 1). In those eight pairs of different species analysed, dDDH values recorded with little variation in the phenotypic properties that (Table S26-S33) supported the proposal of heterotypic synonyms or subspecies.

Totally five heterotypic synonyms and two subspecies are proposed within those genera, they are: *Caldimonas taiwanensis* as the later heterotypic synonym of *Caldimonas manganoxidans*; *Cronobacter dublinensis* subsp. *lactaridi* as the later heterotypic synonym of *Cronobacter dublinensis* subsp. *lausannensis*; *Methylomicrobium agile* as the later heterotypic synonym of *Methylomicrobium album, Paraglaciecola agarilytica* as the later heterotypic synonym of *Paraglaciecola chathamensis* and *Tepidiphilus thermophilus* as the later heterotypic synonym of *Tepidiphilus succinatimandens.* The different subspecies proposed includes *Desulfotignum balticum* subsp. *phosphitoxidans* subsp. nov. and *Haemophilus influenzae* subsp. *aegyptius* subsp. nov. Analysis of the *Neisseria* clade showed the presence of two subspecies as the dDDH value recorded (Table 1) was within the threshold (70-79 %) proposed for subspecies delineation [18], for these two subspecies, similarity was also noticed in the substrate assimilation pattern and enzyme activity (Table S34-35). The subspecies proposed are *Neisseria sicca* subsp. *macacae* subsp. nov. and *Neisseria mucosa* subsp. *cerebrosus* subsp. nov.

Lastly, two important *Proteobacteria* clade comprised of *Pseudoalteromonas* and *Shewanella* were anlaysed for heterotypic synonyms. The *Pseudoalteromonas* clade comprised of *Pseudoalteromonas agarivorans, Pseudoalteromonas donghaensis, Pseudoalteromonas issachenkonii, Pseudoalteromonas atlantica, Pseudoalteromonas lipolytica* and *Pseudoalteromonas tetraodonis* (Fig. 6) all strains sharing high 16S rRNA gene simlairty (Fig. 6 & Table 1) When these species were subjected for dDDH analysis, results supported that among each pair of different species identified in this clade, one may be a heterotypic synonym owing to their strain’s genome similarity in the form of OGRI metrics (Table 1), they were also critically examined for their phenotypic traits wherein, their similarity in phenotypic and biochemical traits are noticed (Table S36-S38). When the 16S rRNA gene sequence-based phylogeny (Fig. 6) was carefully examined, two other taxa of *Pseudoalteromonas* was also phylogenetically close. When they were subjected to OGRI analysis, *Pseudoalteromonas espejiana* showed < 70 dDDH with any other *Pseudoalteromonas* in the clade and for the other *Pseudoalteromonas* species, currently genome data was unavailable. Based on these polyphasic evidences *Pseudoalteromonas agarivorans* was proposed as later heterotypic synonym of *Pseudoalteromonas atlantica*. *Pseudoalteromonas donghaensis* as the later heterotypic synonym of *Pseudoalteromonas lipolytica* and *Pseudoalteromonas issachenkonii* as the later heterotypic synonym of *Pseudoalteromonas tetraodonis*.

The clade comprised of *Shewanella* found to contain two pairs of species (*Shewanella algae* and *Shewanella upenei*; *Shewanella japonica* and *Shewanella pacifica*). When their dDDH was analysed, closeness in their 4 strains genome was observed (Table 1), phenotypic data also supported the fact that between these two pairs most of the biochemical properties are shared (Table S39-S40). These polyphasic evidences support the proposal of *Shewanella upenei* as later heterotypic synonym of *Shewanella algae* and *Shewanella pacifica* as the later heterotypic synonym of *Shewanella japonica*.

### Thermotogae

Genetic closeness in the form of 16S rRNA gene similarity was identified in two genera that belonged to the phylum *Thermotogae* (Fig. 7). A species pair of *Pseudothermotoga* (*P. elfii* and *P. lettingae*) and *Thermotoga* pair (*T. petrophila* and *T. naphthophila*) were selected based 16S rRNA gene similarity, they were subjected to dDDH analysis (Table 1). High similarity in their phenotypic and biochemical traits are observed (Table S41 - S42). The 16S rRNA gene based phylogenetic analysis combined with OGRI metrics and biochemical characters supported the proposal of *Pseudothermotoga lettingae* as the later heterotypic synonym of *Pseudothermotoga elfii*. Whereas in another pair, *T. naphthophila* was hierarchally reduced to subspecies *Thermotoga petrophila* subsp. *naphthophila* subsp. nov.

**Fig. 7.**
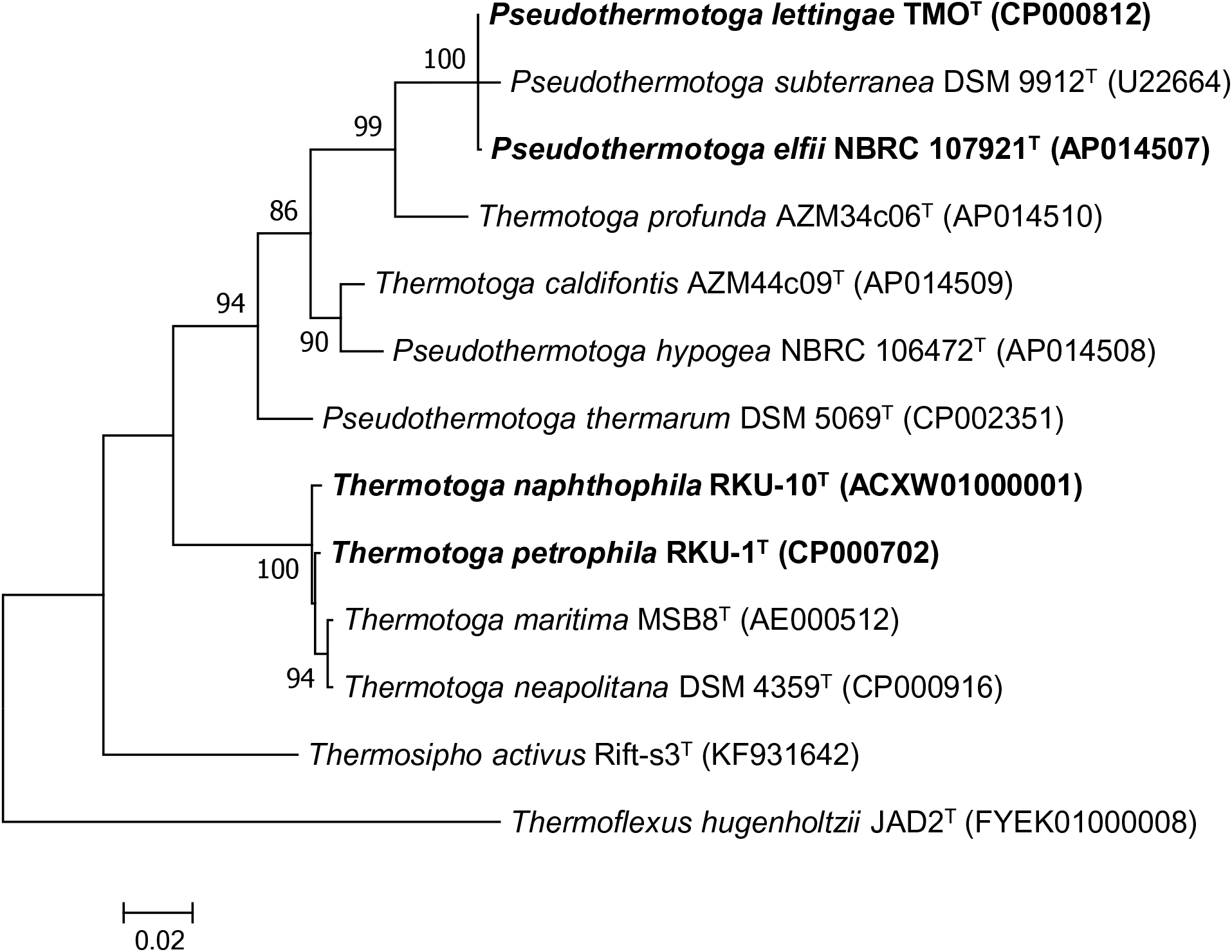
16S rRNA gene sequence-based Maximum likelihood phylogenetic tree showing four heterotypic synonyms from two different genera (in bold) and the closest relatives with validly published names that belonged to phylum *Thermotogae*. Bootstrap values are expressed as percentages of 1000 replications and are shown at branch nodes (>70 %). GenBank accession numbers are indicated in parentheses. Heterotypic synonyms of the type species are marked in bold. Scale bar, 0.02 nt substitutions per position.

### Taxonomic consequences: New (combinations for) species

When the 16S rRNA gene similarity between the species of the different families or phylum were examined, *Firmicutes* and *Proteobacteria* members namely *Saccharococcus thermophilus* and *Haemophilus piscium* were taxonomically incorrect in their genus level placement.

*Saccharococcus thermophilus* was isolated from native sugar beet extraction plants in Sweden. Based on cell morphology and its inability to form endospores, it could be differentiated from the *Geobacillus*, and this trait was also considered as an important criterian in creation of this genus [41]. In the present study, when the 16S rRNA gene phylogeny of this species was analysed, it found to cluster within *Parageobacillus*, for instance, based on 16S rRNA gene similarity (98.9 %), it was phylogenetically similar to *Parageobacillus caldoxylosilyticus*, further the high ANI value (90.3 %) recorded with *P. caldoxylosilyticus* also supported the claim that it belongs to *Parageobacillus* species, since ANI more than 95 % was regarded a OGRI criteria supporting species similarity [23]. Thus, based on the 16S rRNA gene similarity and the tree topology in combination with the OGRI metrics strongly supported the proposal of *Saccharococcus thermophilus* as *Parageobacillus thermophilus* as combination novel. Interestingly, the only other member of *Saccharococcus* was *S. caldoxylosilyticus*, an obligately thermophilic, xylose utilizing, endospore forming bacteria [42] which was subsequently reclassified to *Parageobacillus* as a heterotypic synonym [43] and a recent phylogenomics based reassessment study also insist the presence of such a heterotypic synonym in *Geobacillus* [33].

The *Proteobacteria* genus *H. piscium* was genetically distant from *Haemophilus* genus, its 16S rRNA gene sequence similarity with the *Haemophilus* type species (*H influenzae*) was 85.6 %. In the 16S rRNA gene sequence based analyis it was genetically close to *Aeromonas salmonicida* subsp. *salmonicida* and *Aeromonas salmonicida* subsp. *smithia* with 99.9 and 99.6 % similarity. However, the genome data was currently unavailable, hindering the OGRI analysis. Based on the 16S rRNA tree topology and on the similarity of the *H. piscium* 16S rRNA gene sequence with the *Aeromonas salmonicida* subsp. *salmonicida* and its less similiar with the *Haemophilus* type species, *Haemophilus piscium* was reclassified as *Aeromonas salmonicida* subsp. *piscium*.

#### TAXONOMIC CONSEQUENCES: EMENDATION OF SPECIES AND SUBSPECIES ACTINOBACTERIA

##### EMENDED DESCRIPTION OF *ACTINOKINEOSPORA MZABENSIS* AOUICHE *ET AL.* 2015

Heterotypic synonym: *Actinokineospora spheciospongiae* Kämpfer *et al.* 2015.

The species description is as given before [44, 45] with the following additions. The colonies exhibited variable growth in terms of aerial (pinkish to white) and substrate mycelial (purple to blackish and yellow to tan) pigments. Strains used cellulose, xylose, fructose, maltose and mannitol as sole carbon source, they also reduced nitrate and liquified gelatin. Variability was noticed in utilization of galactose, sucrose and rhamnose as sole carbon sources. The G+C content of the strains are 72.8 % and the genome size is 7.55 Mbp. The type strain of *Actinokineospora mzabensis* is PAL84^T^ (=CECT 8578^T^ = DSM 45961^T^) with GenBank accession number for the whole-genome sequence is GCA_003182415.1.

##### EMENDED DESCRIPTION OF *DIETZIA CINNAMEA* YASSIN *ET AL.* 2006

Heterotypic synonym: *Dietzia papillomatosis* Jones *et al.* 2008.

The species description is as given before [46, 47] with the following additions. The colonies produced orange colour, strains grew in 7 and 8 % of NaCl (w/v), and strains produced acid using d-fructose as carbon source and utilized L-cysteine as sole carbon and nitrogen source. Variability was noticed in degradation of chitin, L-tyrosine, acid production from d-glucose and d-mannose and sucrose, variability was also noticed in utilization of L-arginine as sole carbon and nitrogen source. The G+C content of the type strain genome is 70.8 % and the approximate genome size is 3.60 Mbp. The GenBank accession number for the whole-genome sequence is GCA_001571065.1. The type strain is IMMIB RIV-399^T^ (=CCUG 50875^T^ = DSM 44904^T^ = JCM 13663^T^ = NBRC 102147^T^).

##### EMENDED DESCRIPTION OF *RHODOCOCCUS OPACUS* KLATTE *ET AL.* 1995

Heterotypic synonym: *Rhodococcus imtechensis* Ghosh *et al.* 2006. The species description is as given before [48, 49] with the following additions. All strains utilized maltose and lactose as sole carbon source and L-serine as sole nitrogen source. Variability among the strains are observed for using of L-arabitol, D-melezitose, D-raffinose, L-rhamnose, 2,4 dinitrophenol and *p*-nitrophenol as sole carbon source and energy and L-serine as sole nitrogen source. Variability among the strains are also observed in Biolog GP2 plate assay for utilization of acetic acid, 2,3 butanediol, α-cyclodextrin, L-fucose, D-galacturonic acid, D-lactic acid methyl ester, lactulose, methyl *α*-D-mannoside, propionic acid, putrescine, L-pyroglutamic acid, D-trehalose and uridine 5-monophosphate as sole carbon source. The G+C content of the type strain genome is 67.3 % and the approximate genome size is 8.53 Mbp. The GenBank accession number for the whole-genome sequence is GCA_001646735.1. The type strain is ATCC 51881^T^ (=CIP 104549^T^ = DSM 43205^T^ = IFO 16217^T^ = JCM 9703^T^ = NBRC 100624^T^ = NBRC 16217^T^).

#### ARCHAEA

##### EMENDED DESCRIPTION OF *HALOFERAX VOLCANII* (MULLAKHANBHAI AND LARSEN 1975) TORREBLANCA *ET AL.* 1986

Heterotypic synonym: *Haloferax lucentense* corrig. Gutierrez *et al.* 2004; *Haloferax alexandrinus* Asker and Ohta 2002.

The species description is as given before [50-53] with the following additions. Cell are Gram negative, producing red to pinkish colonies, growth is observed under an optimum temperature of 37 °C, pH of 7.0 and NaCl concentration of 1-4.5 % w/v. All strains are oxidase positive and able to produce H_2_S from thiosulfate and produced acid from arabinose, and unable to hydrolyze starch and casein but variability observed in gelatin hydrolysis. Other variable features observed are cell motility, nitrate reduction, and usage of sucrose as sole carbon source. The G+C content of the type strain genome is 65.5 % and the approximate genome size is 4.01 Mbp. The GenBank accession number for the whole-genome sequence is GCA_000025685.1. The type strain is DS2^T^ (=ATCC 29605^T^ = DSM 3757^T^ = IFO 14742^T^ = JCM 8879^T^ = NBRC 14742^T^ = NCCB 85050^T^ = NCIMB 2012^T^ = VKM B-1768^T^).

##### EMENDED DESCRIPTION OF *METHANOBACTERIUM VETERUM* KRIVUSHIN *ET AL.* 2010

Heterotypic synonym: *Methanobacterium arcticum* Shcherbakova *et al.* 2011.

The species description is as given before [54, 55] with the following additions. Cells are Gram negative, rod shaped with cyst like appearance, strains exhibit growth in temperature range of 15-45 ^ο^C, pH of 5.5 – 8.5 and NaCl concentration of 0-3 % (w/v). Variability was observed in utilization of substrates such as formate, methylamine/H_2_ and methanol/H_2_ and stimulation of growth in presence of acetate. The G+C content of the type strain genome is 33.2 % and the approximate genome size is 3.37 Mbp. The GenBank accession number for the whole-genome sequence is GCA_000745485.1. The type strain is MK4^T^ (=DSM 19849^T^= VKM B-2440^T^).

##### EMENDED DESCRIPTION OF *METHANOSARCINA MAZEI* CORRIG. (BARKER 1936) MAH AND KUHN 1984

Heterotypic synonym: *Methanosarcina soligelidi* Wagner *et al.* 2013.

The species description is as given before [56, 57, 58] with the following additions. Cell are Gram negative, irregular cocci, exhibiting wider temperature range of 0-50 ^ο^C and narrow pH 6.1-8.0 for growth. Optimum NaCl concentration of 0.02 – 0.3 % (w/v) supported the growth of the strains. Strains utilized H_2_/CO_2_, methanol, acetate. Strains cell membrane possessed archaeol phosphatidylglycerol, hydroxyarchaeol phosphatidylglycerol, archaeol phosphatidylethanolamine and hydroxyarchaeol phosphatidylethanolamine as predominant lipids. Variability among strains was noticed in the utilization of dimethylamine. The G+C content of the type strain genome is 41.4 % and the approximate genome size is 4.14 Mbp. The GenBank accession number for the whole-genome sequence is GCA_000970205.1. The type strain is OCM 26^T^ (=DSM 2053^T^ = DSM 3318^T^ = S-6^T^ =VKM B-1636^T^).

#### BACTEROIDETES

##### EMENDED DESCRIPTION OF *RUFIBACTER RUBER* KÝROVÁ *ET AL.* 2016

Heterotypic synonym: *Rufibacter quisquiliarum* Felföldi *et al.* 2016.

The species description is as given before [59, 60] with the following additions. Cells are Gram strain negative, producing pinkish to red coloured colonies on R2A medium, strains positive for catalase, requires a temperature range of 20-37 °C, pH – 7-11 and NaCl concentration range of 0-1 % (w/v) for growth. Strains able to assimilate d-mannose, N-acetylglucosamine, aesculin, glycogen, gentiobiose, *α*-D-glucose, maltose and D-trehalose. All strains produced esterase lipase (C8), leucine arylamidase, valine arylamidase activity and α-Glucosidase enzyme. Difference in assimilation of d-galactose, cellobiose, lactose, melibiose, sucrose and raffinose was observed. The G+C content of the type strain genome is 51.5 % and the approximate genome size is 5.5 Mbp. The GenBank accession number for the whole-genome sequence is GCA_001647275.1. The type strain is CCM 8646^T^ (=LMG 29438^T^).

#### FIRMICUTES

##### EMENDED DESCRIPTION OF CALDANAEROBACTER SUBTERRANEUS SUBSP. TENGCONGENSIS (XUE ET AL. 2001) FARDEAU ET AL. 2004

Heterotypic synonym: *Caldanaerobacter subterraneus* subsp. *yonseiensis* (Kim *et al.* 2001) Fardeau *et al.* 2004.

The species description is as given before [60, 61, 62, 63] with the following additions. Cells grows in a temperature range of 50-80 ^ο^C and pH range of 4.5 - 9 and NaCl concentration of 0-2.5 % (w/v), strains assimilate CO, produced L-alanine, acetate, H_2_ and CO_2_ as diagnostic fermentation products from glucose. Variability was observed in production of lactate as a diagnostic fermentation product from glucose. The G+C content of the type strain genome is 37.6 % and the approximate genome size is 2.69 Mbp. The GenBank accession number for the whole-genome sequence is GCA_000007085.1. The type strain is MB4^T^ (=Chinese Collection of Microorganisms AS 1.2430^T^ = DSM 15242^T^ = JCM 11007^T^ = NBRC 100824^T^).

##### EMENDED DESCRIPTION OF *CARNOBACTERIUM INHIBENS* SUBSP. *INHIBENS* (JÖBORN *ET AL.* 1999) NICHOLSON *ET AL.* 2015

Heterotypic synonym: *Carnobacterium inhibens* subsp. *gilichinskyi* Nicholson *et al.* 2015.

The species description is as given before [64, 65] with the following additions. Cells grows at a wider temperature and pH range of 0-37 ^ο^C and 5.5-9.0 respectively. Cells utilized maltose, D- Mannose, sucrose and trehalose as substrate. Variability was observed in utilization of substrates such as d-galactose, gentiobiose, d-glucose, pectin, potassium tellurite, starch and tetrazolium blue. The G+C content of the type strain genome is 34.9 % and the approximate genome size is 2.75 Mbp. The GenBank accession number for the whole-genome sequence is GCA_000746825.1. The type strain is K1^T^ (=CCUG 31728^T^= CIP 106863^T^= DSM 13024^T^= JCM 16168^T^).

##### EMENDED DESCRIPTION OF *GEOBACILLUS THERMOLEOVORANS* (ZARILLA AND PERRY 1988) NAZINA *ET AL.* 2001

Heterotypic synonym: *Geobacillus kaustophilus* (Priest *et al.* 1989) Nazina *et al.* 2001.

The species description is as given before [31, 66, 67] with the following additions. Strains are positive for catalase activity and nitrate reduction, production of acids from glucose, fructose and maltose and mannose, hydrolysis of gelatin and starch. However, variability was observed among the strains in acid production from adonitol, cellobiose, glycerol, mannitol sucrose, trehalose and D-xylose. Variability was also observed in aesculin and casein hydrolysis, utilization of lactate and citrate as growth substrates. The G+C content of the type strain genome is 52.3 % and the approximate genome size is 3.5 Mbp. The GenBank accession number for the whole-genome sequence is GCA_001610955.1. The type strain is 2 LEH-1^T^ (=ATCC 43513^T^ = BGSC 96A1^T^ = DSM 5366^T^ = LMG 9823^T^).

##### EMENDED DESCRIPTION OF *PARAGEOBACILLUS TOEBII* (SUNG *ET AL.* 2002) ALIYU *ET AL.* 2019

Heterotypic synonym: *Geobacillus galactosidasius* Poli *et al.* 2012; *Geobacillus yumthangensis* Najar *et al.* 2018.

The species description is as given before [33, 68, 69, 70] with the following additions. Abiotic factors (temperature, pH and NaCl concentration) supporting the culture growth was similar for all the strains, however, difference in the biochemical properties such as acid formation form cellobiose, galactose, ribose, glycerol, lactose and rhamnose was observed, additionally variation was observed in the carbon utilization profile. The G+C content of the type strain genome is 42.4 % and the approximate genome size is 3.32 Mbp. The GenBank accession number for the whole- genome sequence is GCA_003688615.1. The type strain is BK-1^T^ (=DSM 14590^T^ = KCTC 0306BP^T^ = LMG 23037^T^ = NBRC 107807^T^ = SK-1^T^).

##### EMENDED DESCRIPTION OF *MEGAMONAS FUNIFORMIS* SAKON *ET AL.* 2008

Heterotypic synonym: *Megamonas rupellensis* Chevrot *et al.* 2008.

The species description is as given before [71, 72] with the following additions. Strains produced acids from glucose, maltose, lactose, mannose, arabinose, salicin, xylose and raffinose. Variability was noticed in acid production from cellobiose and glycerol. The G+C content of the type strain genome is 31.5 % and the approximate genome size is 2.57 Mbp. The GenBank accession number for the whole-genome sequence is GCA_010669225.1. The type strain is YIT 11815^T^ (=DSM 19343^T^ = JCM 14723^T^).

##### EMENDED DESCRIPTION OF THERMOANAEROBACTER THERMOCOPRIAE (JIN ET AL. 1988) COLLINS ET AL. 1994

Heterotypic synonyms: *Thermoanaerobacter italicus* Kozianowski *et al.* 1998; *Thermoanaerobacter mathranii* subsp. *mathranii* (Larsen *et al.* 1998) Carlier *et al.* 2007.

The species description is as given before [73-77] with the following additions. Cells exhibiting Gram positive to variable reaction and non-spore forming, requires an optimum temperature of 60 ^ο^C for growth. All strains degrade xylan, strains utilized glucose as growth substrate and fermentation products are produced using acetate, lactate and ethanol. Variability was noticed in motility and utiliztaio nof sucrose and xylose as growth substrates. The G+C content of the type strain genome is 34.1 % and the approximate genome size is 2.46 Mbp. The GenBank accession number for the whole-genome sequence is GCA_000518565.1. The type strain is JT3-3^T^ (=ATCC 51646^T^= IAM 13577^T^= JCM 7501^T^).

##### EMENDED DESCRIPTION OF *THERMOANAEROBACTER BROCKII* SUBSP. *FINNII* (SCHMID *ET AL.* 1986) CAYOL *ET AL.* 1995

Heterotypic synonym: *Thermoanaerobacter pseudethanolicus* Onyenwoke *et al.* 2007.

The species description is as given before [78-80] with the following additions. Cells exhibit Gram variable reaction and produced spores and motile. Strains used glucose, sucrose, xylose as growth substrates and fermentation products obtained from ethanol. The G+C content of the type strain genome is 34.5 % and the approximate genome size is 2.34 Mbp. The GenBank accession number for the whole-genome sequence is GCA_000175295.2. The type strain is Ako-1^T^ (=ATCC 43586^T^= DSM 3389^T^).

##### EMENDED DESCRIPTION OF *THERMOANAEROBACTER ETHANOLICUS* WIEGEL AND LJUNGDAHL 1982

Heterotypic synonyms: *Thermoanaerobacter wiegelii* Cook *et al.* 1996; *Thermoanaerobacter siderophilus* Slobodkin *et al.* 1999.

The species description is as given before [81-83] with the following additions. Cells exhibit Gram variable reaction with variability noticed in spore production. All strains are motile, capable of starch hydrolysis, used glucose, sucrose, xylose as substrate for growth and fermentation products are formed using acetate, lactase and ethanol as carbon sources. Variability among the strains are noticed in utilization of ribose, mannitol, glycerol, pyruvate as growth substrates. The G+C content of the type strain genome is 34.2 % and the approximate genome size is 2.91 Mbp. The GenBank accession number for the whole-genome sequence is GCA_003722315.1. The type strain is JW 200^T^ (=ATCC 31550^T^= DSM 2246^T^).

#### NITROSPIRAE

##### EMENDED DESCRIPTION OF THERMODESULFOVIBRIO YELLOWSTONII HENRY ET AL. 1994

Heterotypic synonym: *Thermodesulfovibrio islandicus* Sonne-Hansen and Ahring 2000.

The species description is as given before [35-36] with the following additions. Nutritionally strains can utilize pyruvate, hydrogen (plus acetate) and formate (plus acetate) as both electron donor and carbon source in the presence of sulfate as terminal electron acceptor and in addition all strains utilized sulfate and thiosulfate as electron acceptors. However, variability was noticed in utilization of sulfite and nitrate as terminal electron acceptors. Strains showed fermentative growth with pyruvate. Strains preferred a thermophilic range of temperature (40 - 70 ^ο^C) for growth. The G+C content of the type strain genome is 34.1 % and the approximate genome size is 2.0 Mbp. The GenBank accession number for the whole-genome sequence is GCA_000020985.1. The type strain is YP87^T^ (=ATCC 51303^T^= DSM 11347^T^).

#### PROTEOBACTERIA

##### EMENDED DESCRIPTION OF *AEROMONAS SALMONICIDA* (LEHMANN AND NEUMANN 1896) GRIFFIN *ET AL.* 1953 (APPROVED LISTS 1980)

Heterotypic synonym: *Aeromonas salmonicida* subsp. *masoucida* Kimura 1969 (Approved Lists 1980).

The species description is as given before [85, 86, 87] with the following additions. Motile cells utilized urocanic acid, sucrose and fermented mannitol. However, variability noticed in brown diffusible pigment production, indole production and Voges-Proskauer test at 25 ^ο^C, fermentation of sucrose and gas production from glucose. The G+C content of the type strain genome is 58.3 % and the approximate genome size is 4.94 Mbp. The GenBank accession number for the whole-genome sequence is GCA_900445115.1. The type strain is ATCC 33658^T^ (=CAIM 346^T^ = CIP 103209^T^ = DSM 18220^T^ = DSM 19634^T^ = HAMBI 1984^T^ = JCM 7874^T^ = LMG 3780^T^ = NCIMB 1102^T^ = NCTC 12959^T^).

##### EMENDED DESCRIPTION OF *ALTEROMONAS STELLIPOLARIS* VAN TRAPPEN *ET AL.* 2004

Heterotypic synonym: *Alteromonas addita* Ivanova *et al.* 2005.

The species description is as given before [37, 38] with the following additions. All strains are Gram-negative, motile, oxidase- and catalase-positive and negative for indole and H_2_S production, grow at 3–6 % NaCl and produce lipase (Tween 80). Strains able to grow at 4 ^ο^C but growth was not observed at 40 ^ο^C. Similarly, able to grow at 10 % but growth was not observed at 15 % NaCl. Strains hydrolyzed starch. Variability observed in haemolysis and assimilation of D-mannitol and L-lactate. The G+C content of the type strain genome is 43.5 % and the approximate genome size is 4.90 Mbp. The GenBank accession number for the whole-genome sequence is GCA_001562115.1. The type strain is ANT 69a^T^ (=DSM 15691^T^ = LMG 21861^T^).

##### EMENDED DESCRIPTION OF *BORDETELLA BRONCHISEPTICA* (FERRY 1912) MORENO-LÓPEZ 1952 (APPROVED LISTS 1980)

Heterotypic synonym: *Bordetella parapertussis* (Eldering and Kendrick 1938) Moreno-López 1952 (Approved Lists 1980); *Bordetella pertussis* (Bergey *et al.* 1923) Moreno-López 1952 (Approved Lists 1980). The species description is as given before [88-91] with the following additions. Minute coccobacillus motile or non-motile cells, variability in oxidase activity and growth on the MacConkey agar and Simmons citrate agar was noticed. Differences in urease activity, reduction of nitrate to nitrite, brown pigment production on HI agar with tyrosine, litmus milk alkalization was observed among the strains. None of the strains assimilated D-xylose and D-gluconate but variability observed among assimilation of phenyl acetate. All strains are negative for lipase (C14) and valine arylamidase, but variability observed on chymotrypsin and naphthol-AS-Bi- phosphohydrolase activity. The G+C content of the type strain genome is 68.2 % and the approximate genome size is 5.17 Mbp. The GenBank accession number for the whole-genome sequence is GCA_900445725.1. The type strain is ATCC 19395^T^ (=CCUG 219^T^ = CIP 55.110^T^ = DSM 13414^T^ = IFO 13691^T^ = LMG 1232^T^ = NBRC 13691^T^ = NCTC 452^T^).

##### EMENDED DESCRIPTION OF *CALDIMONAS MANGANOXIDANS* TAKEDA *ET AL.* 2002

Heterotypic synonym: *Caldimonas taiwanensis* Chen *et al.* 2005.

The species description is as given before [92, 93] with the following additions. Gram negative rods, for growth, strains require an optimum temperature and pH of 50 ^ο^C and 7.0 respectively. Strains exhibit variable oxidase activity and variable utilization of substrates like galactose, malate, malonate, sucrose, acetate, fructose and trehalose. The G+C content of the type strain genome is 66 % and the approximate genome size is 3.53 Mbp. The GenBank accession number for the whole-genome sequence is GCA_000381125.1. The type strain is HS^T^ (=ATCC BAA- 369^T^ = IFO 16448^T^= JCM 10698^T^= NBRC 16448^T^).

##### EMENDED DESCRIPTION OF CRONOBACTER DUBLINENSIS SUBSP. LAUSANNENSIS IVERSEN ET AL. 2008

Heterotypic synonym: *Cronobacter dublinensis* subsp. *lactaridi* Iversen *et al.* 2008.

The species description is as given before [95] with the following additions. Strains variable Indole production, they utilized *cis*-Aconitate, *trans*-aconitate, palatinose, 1-O-methyl-*α*- glucopyranoside and 4-aminobutyrate. But variability observed in the utilization of lactulose, maltitol, putrescine, turanose, *myo*-inositol. However, strains area unable to utilize dulcitol, malonate and Melezitose, The G+C content of the type strain genome is 57.9 % and the approximate genome size is 4.61 Mbp. The GenBank accession number for the whole-genome sequence is GCA_000409365.1. The type strain is E515^T^ (=DSM 18706^T^= JCM 16469^T^ = LMG 23824^T^).

##### EMENDED DESCRIPTION OF *METHYLOMICROBIUM ALBUM* (BOWMAN *ET AL.* 1993) BOWMAN *ET AL.* 1995

Heterotypic synonym: *Methylomicrobium agile* (Bowman *et al.* 1993) Bowman *et al.* 1995.

The species description is as given before [95, 96] with the following additions. Cells are motile, non-cyst forms able to grow at an optimum temperature of 25-30 ^ο^C but unable to grow at 3 % NaCl concentration and the optimum pH observed for growth is 7.0. Strains possessed particulate methane mono oxygenase, but soluble methane mono oxygenase is absent. Strain level variability was observed during growth at 37 ^ο^C. The G+C content of the type strain genome is 56.2 % and the approximate genome size is 4.49 Mbp. The GenBank accession number for the whole-genome sequence is GCA_000214275.3. The type strain is BG8^T^ (=ACM 3314^T^ = ATCC 33003^T^ = NCIMB 11123^T^ = VKM-BG8^T^).

##### EMENDED DESCRIPTION OF *PARAGLACIECOLA CHATHAMENSIS* (MATSUYAMA *ET AL.* 2006) SHIVAJI AND REDDY 2014

Heterotypic synonym: *Paraglaciecola agarilytica* (Yong *et al.* 2007) Shivaji and Reddy 2014.

The species description is as given before [97, 98, 99] with the following additions. Strains were able to grow at temperature range of 7-30 ^ο^C but growth was variable at 4 ^ο^C, the NaCl range of 2-8 % supported the growth. Strains hydrolyzed aesculin, casein and starch; utilized D-galactose, D-mannitol, D-mannose as carbon source. Strain variability was observed in usage of acetate, cellobiose, maltose and sucrose as carbon substrates and hydrolysis of ONPG. The G+C content of the type strain genome is 44.1 % and the approximate genome size is 5.27 Mbp. The GenBank accession number for the whole-genome sequence is GCA_000314955.1. The type strain is S18K6^T^ (=JCM 13645^T^= NCIMB 14146^T^).

##### EMENDED DESCRIPTION OF *PSEUDOALTEROMONAS ATLANTICA* (AKAGAWA- MATSUSHITA *ET AL.* 1992) GAUTHIER *ET AL.* 1995

Heterotypic synonym: *Pseudoalteromonas agarivorans* Romanenko *et al.* 2003.

The species description is as given before [39, 100, 101] with the following additions. Strains are motile, positive for the sodium-ion requirement for growth, growth at 25–28 °C, produced acid from mannitol and oxidized glycogen. Variability was noticed in hydrolysis of carrageenan, acid production from glucose, D-mannitol, maltose, caprate, phenylacetate. Strain level variation was also observed in oxidation of D-fructose, mannose, sucrose, glycerol, citrate, propionate butyrate. The G+C content of the type strain genome is 40.8 % and the approximate genome size is 4.47 Mbp. The GenBank accession number for the whole-genome sequence is GCA_007988745.1. The type strain is NCIMB 301^T^ (=ATCC 19262^T^ = CIP 104721^T^ = DSM 6840^T^ = IAM 12927^T^ = JCM 8845^T^ = NBRC 103033^T^).

##### EMENDED DESCRIPTION OF PSEUDOALTEROMONAS LIPOLYTICA XU ET AL. 2010

Heterotypic synonym: *Pseudoalteromonas donghaensis* Oh *et al.* 2011.

The species description is as given before [102, 103] with the following additions. Strains are positive for catalase and oxidase activity, nitrate reduction, hydrolyzed gelatin and Tween 80, they utilized N-acetyl-glucosamine, L-arabinose, maltose and mannose as sole carbon source. Acid is produced from maltose and mannose as substrates. Variability among strains are observed in acid production from glucose. The G+C content of the type strain genome is 41.4 % and the approximate genome size is 4.54 Mbp. The GenBank accession number for the whole-genome sequence is GCA_900116435.1. The type strain is LMEB 39^T^ (=CGMCC 1.8499^T^= DSM 22356^T^ = JCM 15903^T^).

##### EMENDED DESCRIPTION OF *PSEUDOALTEROMONAS TETRAODONIS* (SIMIDU *ET AL.* 1990) IVANOVA *ET AL.* 2001

Heterotypic synonym: *Pseudoalteromonas issachenkonii* Ivanova *et al.* 2002.

The species description is as given before [104-106] with the following additions. Strains grow at 4 ^ο^C but strain variability noticed for growth at 37 ^ο^C, strain growth is noticed upto 12 % NaCl concentration. DNase activity observed in strains, but variability noticed in Chitinase activity but none of the strains shows amylase, agarose and carrageenase activity. All strains used D-galactose, sucrose, maltose, citrate, acetate, and pyruvate as substrates. But strain variability was noticed in utilization of D-fructose, melibiose, lactose, succinate and mannitol as substrates. The G+C content of the type strain genome is 40.3 % and the approximate genome size is 4.13 Mbp. The GenBank accession number for the whole-genome sequence is GCA_002310835.1. The type strain is GFC^T^ (=ATCC 51193^T^ = CIP 104758^T^ = CIP 107120^T^ = DSM 9166^T^ = IAM 14160^T^ = JCM 21038^T^ = KMM 458^T^ = NBRC 103034^T^).

##### EMENDED DESCRIPTION OF *SHEWANELLA ALGAE* CORRIG. SIMIDU *ET AL.* 1990

Heterotypic synonym: *Shewanella upenei* Kim *et al.* 2012.

The species description is as given before [106, 107] with the following additions. Strains exhibited motility and positive for motility, catalase, oxidase; H_2_S production, nitrate reduction, hydrolysis of casein, DNA, gelatin, tyrosine and Tweens 20, 40, 60, and 80; utilization of D-glucose, acetate, L-malate, pyruvate and succinate; activity of alkaline phosphatase, esterase (C4), esterase lipase (C8), leucine arylamidase (weak), Į-chymotrypsin. Variability among strains are noticed for acid production from D-glucose and D-ribose, acid phosphatase and naphthol-AS-BI-phosphohydrolase activity. The G+C content of the type strain genome is 53 % and the approximate genome size is 4.87 Mbp. The GenBank accession number for the whole- genome sequence is GCA_009183365.1. The type strain is OK-1^T^ (=ATCC 51192^T^= CCUG 39064^T^ = CIP 106454^T^ = DSM 9167^T^ = IAM 14159^T^ = JCM 21037^T^ = NBRC 103173^T^).

##### EMENDED DESCRIPTION OF *SHEWANELLA JAPONICA* IVANOVA *ET AL.* 2001

Heterotypic synonym: *Shewanella pacifica* Ivanova *et al.* 2004.

The species description is as given before [109, 110] with the following additions. Strains are oxidase and catalase positive, reduce nitrate to nitrite, hemolytic and produced lipase, amylase, gelatinase and Chitinase, strains can grow at 32 ^ο^C but variability in growth is noticed at 4 ^ο^C and growth at 6 % NaCl. Strain level difference is also noticed in the utilization of d-galactose and succinate as substrate. The G+C content of the type strain genome is 40.8 % and the approximate genome size is 4.98 Mbp. The GenBank accession number for the whole-genome sequence is GCA_002075795.1. The type strain is ATCC BAA-316^T^ (=CIP 106860^T^ = DSM 15915^T^ = JCM 21433^T^ = KMM 3299^T^ = LMG 19691^T^ = NBRC 103171^T^).

##### EMENDED DESCRIPTION OF *TEPIDIPHILUS SUCCINATIMANDENS* (BONILLA SALINAS *ET AL.* 2004) PODDAR *ET AL.* 2014

Heterotypic synonym: *Tepidiphilus thermophilus* Poddar *et al.* 2014.

The species description is as given before [110, 111] with the following additions. All strains used Tween 40, Tween 80, D-glucuronic acid and glucuronamide as substrates. Variability was noticed in Urea hydrolysis and nitrite reduction, strain level difference is observed for valine arylamidase, cysteine arylamidase, trypsin and *α*-glucosidase activity. Variability in oxidation of carbon sources such as i-erythritol, xylitol, *cis*-aconitic acid, D-galacturonic acid, *α*-ketoglutaric acid, *α*-ketovaleric acid, DL-lactic acid, D-alanine, L-alanine, L-asparagine, L- glutamic acid, L- leucine, L-serine, glycerol and D-glucose 6-phosphate. The G+C content of the type strain genome is 65.9 % and the approximate genome size is 2.37 Mbp. The GenBank accession number for the whole-genome sequence is GCA_006503695.1. The type strain is 4BON ^T^ (= CIP 107790^T^ = DSM 15512^T^).

#### THERMOTOGAE

##### EMENDED DESCRIPTION OF *PSEUDOTHERMOTOGA ELFII* (RAVOT *ET AL.* 1995) BHANDARI AND GUPTA 2014

Heterotypic synonym: *Pseudothermotoga lettingae* (Balk *et al.* 2002) Bhandari and Gupta 2014.

The species description is as given before [112-114] with the following additions. All strains can grow at a temperature range of 50-70 ^ο^C, pH range of 5.5 – 8.5 and NaCl concentration of 1.2 %. Strain utilized xylose a carbon source. However, strain level variability was observed in the reduction of elemental sulfur and usage of acetate+thiosulfate as the substrate for growth. The G+C content of the type strain genome is 38.7 % and the approximate genome size is 2.17 Mbp. The GenBank accession number for the whole-genome sequence is GCA_000504085.1. The type strain is SERB^T^ (=ATCC 51869^T^= DSM 9442^T^ = SEBR 6459^T^).

### Taxonomic Consequences: New Subspecies

#### ACTINOBACTERIA

##### DESCRIPTION OF *NOCARDIA EXALBIDA* SUBSP. *GAMKENSIS* SUBSP. NOV

*Nocardia exalbida* subsp. *gamkensis* (N.L. masc./fem. adj. *gamkensis*, pertaining to the river Gamka in South Africa)

The description is as given for *Nocardia gamkensis* [115]. The G+C content of the type strain genome is 68.4 % and the approximate genome size is 7.71 Mbp. The GenBank accession number for the whole-genome sequence is GCA_001612985.1. The type strain is CZH20^T^ (=DSM 44956^T^= JCM 14299^T^ = NRRL B-24450^T^).

##### DESCRIPTION OF *NOCARDIA IGNORATA* SUBSP. *COUBLEAE* SUBSP. NOV

*Nocardia ignorata* subsp. *coubleae* (N.L. gen. fem. n. *coubleae*, of Couble, named after Andrée Couble, in recognition of her contribution to the French Nocardiosis Observatory, Lyon, France).

The description is as given for *Nocardia coubleae* [116]. The G+C content of the type strain genome is 67.9 % and the approximate genome size is 6.62 Mbp. The GenBank accession number for the whole-genome sequence is GCA_001612805.1. The type strain is OFN N12^T^ (=CIP 108996T= DSM 44960T= JCM 15318^T^).

##### DESCRIPTION OF *NOCARDIA NOVA* SUBSP. *ELEGANS* SUBSP. NOV

*Nocardia nova* subsp. *elegans* (e’le.gans L. adj. *elegans*, fastidious (with respect of utilization of nutrients)

The description is as given for *Nocardia elegans* [117]. The G+C content of the type strain genome is 67.9 % and the approximate genome size is 7.54 Mbp. The GenBank accession number for the whole-genome sequence is GCA_001612845.1. The type strain is IMMIB N- 402^T^ (=CCUG 50200^T^= CIP 108553^T^= DSM 44890^T^= JCM 13374^T^).

#### FIRMICUTES

##### DESCRIPTION OF CARBOXYDOTHERMUS HYDROGENOFORMANS SUBSP. *FERRIREDUCENS* SUBSP. NOV

*Carboxydothermus hydrogenoformans* subsp. *ferrireducens* [fer.ri.re.du’cens L. neut. n. *ferrum*, iron; L. pres. part. *reducens*, converting to a different state; N.L. part. adj. *ferrireducens*, reducing (ferric) iron]

The description is as given for *Carboxydothermus ferrireducens* [118-119]. The G+C content of the type strain genome is 41.9 % and the approximate genome size is 2.44 Mbp. The GenBank accession number for the whole-genome sequence is GCA_000427565.1. The type strain is JW/AS-Y7^T^ (=DSM 11255^T^= VKM B-2392^T^).

##### DESCRIPTION OF GEOBACILLUS STEAROTHERMOPHILUS SUBSP. LITUANICUS SUBSP. NOV

*Geobacillus stearothermophilus* subsp. *lituanicus* (li.tu.a’ni.cus M.L. adj. *lituanicus*, of Lithuania, referring to the Lithuanian oilfield from where the type strain was isolated)

The description is as given for *Geobacillus lituanicus* [32]. The G+C content of the type strain genome is 52.1 % and the approximate genome size is 3.5 Mbp. The GenBank accession number for the whole-genome sequence is GCA_002243605.1. The type strain is N-3^T^ (=DSM 15325^T^ = VKM B-2294^T^).

##### DESCRIPTION OF SALIMICROBIUM SALEXIGENS SUBSP. JEOTGALI SUBSP. NOV

*Salimicrobium salexigens* subsp. *jeotgali* (je.ot.ga’li N.L. gen. n. *jeotgali*, of jeotgal, from which the organism was first isolated).

The description is as given for *Salimicrobium jeotgali* [120]. The G+C content of the type strain genome is 46.3 % and the approximate genome size is 2.78 Mbp. The GenBank accession number for the whole-genome sequence is GCA_001685435.3. The type strain is MJ3^T^ (=JCM 19758^T^= KACC 16972^T^).

#### PROTEOBACTERIA

##### DESCRIPTION OF DESULFOTIGNUM BALTICUM SUBSP. PHOSPHITOXIDANS SUBSP. NOV

*Desulfotignum balticum* subsp. *phosphitoxidans* (phos.phit.o’xi.dans N.L. part. adj. *phosphitoxidans*, oxidizing phosphite, referring to its utilization of this unusual electron source).

The description is as given for *Desulfotignum phosphitoxidans* [121]. The G+C content of the type strain genome is 51.3 % and the approximate genome size is 4.50 Mbp. The GenBank accession number for the whole-genome sequence is GCA_000350545.1. The type strain is FiPS-3^T^ (=DSM 13687^T^= OCM 818^T^).

##### DESCRIPTION OF *HAEMOPHILUS INFLUENZAE* SUBSP. *AEGYPTIUS* SUBSP. NOV

*Haemophilus influenzae* subsp. a*egyptius* (L. masc. adj. *aegyptius*, Aegyptian).

The description is as given for *Haemophilus aegyptius* [122, 123]. The G+C content of the type strain genome is 37.1 % and the approximate genome size is 1.96 Mbp. The GenBank accession number for the whole-genome sequence is GCA_000195005.1. The type strain is ATCC 11116^T^ (=CCUG 25716^T^ = CIP 52.129^T^ = DSM 21187^T^ = NCTC 8502^T^).

##### DESCRIPTION OF *NEISSERIA SICCA* SUBSP. *MACACAE* SUBSP. NOV

*Neisseria sicca* subsp. *macacae* (ma.ca’cae N.L. gen. n. *macacae*, of a monkey, referring to the source of the isolate (isolated from the oropharynges of rhesus monkeys); from Portuguese n *macaco*, female monkey; from N.L. n. *Macaca*, the generic name of macaques).

The description is as given for *Neisseria macacae* [124]. The G+C content of the type strain genome is 50 % and the approximate genome size is 2.75 Mbp. The GenBank accession number for the whole-genome sequence is GCA_000220865.1. The type strain is M-740^T^ (=ATCC 33926^T^ = CIP 103346^T^ = DSM 19175^T^).

##### DESCRIPTION OF *NEISSERIA MUCOSA* SUBSP. *CEREBROSUS* SUBSP. NOV

*Neisseria mucosa* subsp. c*erebrosus* (L. masc. adj. *cerebrosus*, having a madness of the brain, hare-brained, hotbrained, passionate, intended to mean pertaining to the brain, the original source of isolation of this organism)

The description is as given for *Morococcus cerebrosus* [125]. The G+C content of the type strain genome is 51.4 % and the approximate genome size is 2.45 Mbp. The GenBank accession number for the whole-genome sequence is GCA_000813705.1. The type strain is UQM 858^T^ (=ATCC 33486^T^= CIP 81.93^T^ = DSM 24335^T^= NCTC 11393^T^).

##### DESCRIPTION OF THERMOTOGA PETROPHILA SUBSP. NAPHTHOPHILA SUBSP. NOV

*Thermotoga petrophila* subsp. *naphthophila* (Gr. n. *naphtha*, naphta, bitumen; N.L. fem. adj. *phila*, friend, loving; from Gr. fem. adj. *philê*, loving; N.L. fem. adj. *naphthophila*, bitumen- loving).

The description is as given for *Thermotoga naphthophila* [126]. The G+C content of the type strain genome is 46.1 % and the approximate genome size is 1.81 Mbp. The GenBank accession number for the whole-genome sequence is GCA_000025105.1. The type strain is RKU-10^T^ (=DSM 13996^T^= JCM 10882^T^).

### Taxonomic Consequences: New (Combinations for) Species

#### DESCRIPTION OF *AEROMONAS SALMONICIDA* SUBSP. *PISCIUM* COMB. NOV

*Aeromonas salmonicida* subsp. *piscium* (pis’ci.um L. gen. pl. n. *piscium*, of fishes).

Basonym: *Haemophilus piscium* Snieszko *et al.* 1950 (Approved Lists 1980).

The description is same as given before [127-133] for *Haemophilus piscium* with the following additions. 16S rRNA phylogeny analyses provided strong evidence for assignment of this species to the genus *Aeromonas*. The type strain is ATCC 10801^T^ (=CCUG 15943= CIP 106116= JCM 7872^T^).

#### DESCRIPTION OF PARAGEOBACILLUS THERMOPHILUS COMB. NOV

*Parageobacillus thermophilus* (ther.mo’phi.lus Gr. fem. n. *thermê*, heat; N.L. adj. *philus -a -um*, friend, loving; from Gr. adj. *philos -ê -on*, loving; N.L. masc. adj. *thermophilus*, heat-loving)

Basonym: *Saccharococcus thermophilus* Nystrand 1984.

The description is same as given before [41] for *Saccharococcus thermophilus* with the following additions. Phylogenomic analyses and 16S rRNA phylogeny provided strong evidence for assignment of this species to the genus *Parageobacillus*. The G+C content of the type-strain genome is 44.88 %, its approximate size 3.15 Mbp, its NCBI GenBank assembly accession GCA_011761475.1. The type strain is 657^T^ (=ATCC 43125^T^= CCM 3586^T^= DSM 4749^T^).

