## Supplemental Table S1_S42 for "Genome based analyses reveals the presence of heterotypic synonyms and subspecies in Bacteria and Archaea"

**Table S1.** Sequences used in this study. Unless noted, all genomes and 16S rRNA gene sequences represent the type strain of the respective species and were downloaded from NCBI (https://www.ncbi.nlm.nih.gov) or EzBioCloud database (https://www.ezbiocloud.net/).

| **Species** | **Strain** | **16S rRNA accession number** | **Genbank accession number** |
|---|---|---|---|
| *Actinokineospora mzabensis* | CECT 8578$^{T}$ | KJ504177 | GCA_003182415.1 |
| *Actinokineospora spheciospongiae* | EG49$^{T}$ | AYXG01000061 | GCA_000564855.1 |
| *Aeromonas salmonicida* subsp. *masoucida* | NBRC 13784$^{T}$ | BAWQ01000150 | GCA_000647955.1 |
| *Aeromonas salmonicida* subsp. *salmonicida* | NCTC 12959$^{T}$ | LSGW01000109 | GCA_900445115.1 |
| *Alteromonas addita* | R10SW13$^{T}$ | CP014322 | GCA_001562195.1 |
| *Alteromonas stellipolaris* | LMG 21861$^{T}$ | CP013926 | GCA_001562115.1 |
| *Bordetella bronchiseptica* | NCTC 452$^{T}$ | U04948 | GCA_900445725.1 |
| *Bordetella parapertussis* | FDAARGOS 177$^{T}$ | LRII01000001 | GCA_001525545.2 |
| *Bordetella pertussis* | 18323$^{T}$ | BX470248 | GCA_000306945.1 |
| *Caldanaerobacter subterraneus* subsp. *tengcongensis* | MB4$^{T}$ | AE008691 | GCA_000007085.1 |
| *Caldanaerobacter subterraneus* subsp. *yonseiensis* | KB-1$^{T}$ | AXDC01000042 | GCA_000473865.1 |
| *Caldimonas manganoxidans* | ATCC BAA-369$^{T}$ | AB008801 | GCA_000381125.1 |
| *Caldimonas taiwanensis* | NBRC 104434$^{T}$ | BCWK01000040 | GCA_001592165.1 |
| *Carboxydothermus ferrireducens* | DSM 11255$^{T}$ | U76363 | GCA_000427565.1 |
| *Carboxydothermus hydrogenoformans* | Z-2901$^{T}$ | CP000141 | GCA_000012865.1 |
| *Carnobacterium inhibens* subsp. *gilichinskyi* | WN1359$^{T}$ | CP006812 | GCA_000493735.1 |
| *Carnobacterium inhibens* subsp. *inhibens* | DSM 13024$^{T}$ | JQIV01000006 | GCA_000746825.1 |
| *Corynebacterium afermentans* subsp. *afermentans* | DSM 44280$^{T}$ | jgi.1096551 | GCA_900156035.1 |
| *Corynebacterium afermentans* subsp. *lipophilum* | HSID17239 | X82055 | GCA_003989555.1 |
| *"Corynebacterium ihumii"* * | GD7$^{T}$ | HG001324 | GCA_000403725.1 |
| *Cronobacter dublinensis* subsp. *lactaridi* | LMG 23825$^{T}$ | AJKX01000065 | GCA_000409345.1 |
| *Cronobacter dublinensis* subsp. *lausannensis* | LMG 23824$^{T}$ | AJKY01000076 | GCA_000409365.1 |
| *Desulfotignum balticum* | DSM 7044$^{T}$ | ATWO01000001 | GCA_000421285.1 |
| *Desulfotignum phosphitoxidans* | DSM 13687$^{T}$ | APJX01000002 | GCA_000350545.1 |
| *Dietzia cinnamea* | NBRC 102147$^{T}$ | AJ920289 | GCA_001571065.1 |
| *Dietzia maris* | 97$^{T}$ | X79290 | GCA_004338615.1 |
| *Dietzia papillomatosis* | NBRC 105045$^{T}$ | BCSL01000097 | GCA_001570845.1 |
| *Geobacillus galactosidasius* | DSM 18751$^{T}$ | AM408559 | GCA_002217735.1 |
| *Geobacillus kaustophilus* | NBRC 102445$^{T}$ | BBJV01000091 | GCA_005160065.1 |
| *Geobacillus lituanicus* | N-3$^{T}$ | CP017692 | GCA_002243605.1 |
| *Geobacillus stearothermophilus* | ATCC 12980$^{T}$ | AB271757 | GCA_001277805.1 |
| *Geobacillus thermoleovorans* | KCTC 3570$^{T}$ | CP014335 | GCA_001610955.1 |
| *Geobacillus yumthangensis* | AYN2$^{T}$ | MG603320 | GCA_002494375.1 |
| *Haemophilus aegyptius* | ATCC 11116$^{T}$ | GL878535 | GCA_000195005.1 |
| *Haemophilus influenzae* | NCTC 8143$^{T}$ | LN831035 | GCA_001457655.1 |
| *Haloferax alexandrinus* | JCM 10717$^{T}$ | AB037474 | GCA_000336735.1 |
| *Haloferax lucentense* | DSM 14919$^{T}$ | AOLH01000027 | GCA_000336795.1 |
| *Haloferax volcanii* | DS2$^{T}$ | CP001956 | GCA_000025685.1 |
| *Megamonas funiformis* | YIT 11815$^{T}$ | AB300988 | GCA_010669225.1 |
| *Megamonas rupellensis* | DSM 19944$^{T}$ | EU346729 | GCA_000378365.1 |
| *Methanobacterium arcticum* | M2$^{T}$ | DQ517520 | GCA_000746075.1 |
| *Methanobacterium veterum* | MK4$^{T}$ | EF016285 | GCA_000745485.1 |
| *Methanosarcina mazei* | S-6$^{T}$ | CP009512 | GCA_000970205.1 |
| *Methanosarcina soligelidi* | SMA-21$^{T}$ | JQLR01000001 | GCA_000744315.1 |
| *Methylomicrobium agile* | ATCC 35068$^{T}$ | JPOJ01000001 | GCA_000733855.1 |
| *Methylomicrobium album* | BG8$^{T}$ | CM001475 | GCA_000214275.3 |
| *Morococcus cerebrosus* | CIP 81.93$^{T}$ | JUFZ01000072 | GCA_000813705.1 |
| *Neisseria macacae* | ATCC 33926$^{T}$ | AFQE01000146 | GCA_000220865.1 |

| | | | |
|---|---|---|---|
| *Neisseria mucosa* | ATCC 19696$^{T}$ | AB910739 | GCA_003028315.1 |
| *Neisseria sicca* | ATCC 29256$^{T}$ | ACKO02000016 | GCA_000174655.1 |
| *Nocardia coubleae* | NBRC 108252$^{T}$ | JN041456 | GCA_001612805.1 |
| *Nocardia elegans* | NBRC 108235$^{T}$ | AJ854057 | GCA_001612845.1 |
| *Nocardia exalbida* | NBRC 100660$^{T}$ | BAFZ01000028 | GCA_000308575.1 |
| *Nocardia gamkensis* | NBRC 108242$^{T}$ | JN041479 | GCA_001612985.1 |
| *Nocardia ignorata* | DSM 44496$^{T}$ | BDBI01000064 | GCA_004362495.1 |
| *Nocardia nova* | NBRC 15556$^{T}$ | BDBN01000167 | GCA_001613005.1 |
| *Parageobacillus caldoxylosilyticus* | NBRC 107762$^{T}$ | BAWO01000028 | GCA_000632715.1 |
| *Parageobacillus toebii* | DSM 14590$^{T}$ | BDAQ01000034 | GCA_003688615.1 |
| *Paraglaciecola agarilytica* | NO2$^{T}$ | BAEK01000058 | GCA_000314935.1 |
| *Paraglaciecola chathamensis* | S18K6$^{T}$ | BAEM01000005 | GCA_000314955.1 |
| *Pseudoalteromonas agarivorans* | DSM 14585$^{T}$ | CP011011 | GCA_002310855.1 |
| *Pseudoalteromonas atlantica* | NBRC 103033$^{T}$ | BJUT01000111 | GCA_007988745.1 |
| *Pseudoalteromonas donghaensis* | HJ51$^{T}$ | CP032090 | GCA_003515105.1 |
| *Pseudoalteromonas issachenkonii* | KCTC 12958$^{T}$ | CP013350 | GCA_001455325.1 |
| *Pseudoalteromonas lipolytica* | CGMCC 1.8499$^{T}$ | jgi.1058048 | GCA_900116435.1 |
| *Pseudoalteromonas tetraodonis* | GFC$^{T}$ | CP011041 | GCA_002310835.1 |
| *Pseudothermotoga elfii* | NBRC 107921$^{T}$ | AP014507 | GCA_000504085.1 |
| *Pseudothermotoga lettingae* | TMO$^{T}$ | CP000812 | GCA_000017865.1 |
| *Rhodococcus imtechensis* | RKJ300$^{T}$ | AY525785 | GCA_000260815.1 |
| *Rhodococcus opacus* | DSM 43205$^{T}$ | X80630 | GCA_001646735.1 |
| *Rufibacter quisquiliarum*† | DSM 29854$^{T}$ | KM083132 | 2827981459† |
| *Rufibacter ruber* | CCM 8646$^{T}$ | LRMM01000126 | GCA_001647275.1 |
| *Saccharococcus thermophilu*s | DSM 4749$^{T}$ | X70430 | GCA_011761475.1 |
| *Salimicrobium jeotgali* | MJ3$^{T}$ | AMPQ01000045 | GCA_001685435.3 |
| *Salimicrobium salexigens* | DSM 22782$^{T}$ | jgi.1096523 | GCA_900156705.1 |
| *Shewanella algae* | CECT 5071$^{T}$ | BALO01000089 | GCA_009183365.1 |
| *Shewanella japonica* | KCTC 22435$^{T}$ | CP020472 | GCA_002075795.1 |
| *Shewanella pacifica* | KCTC 12235$^{T}$ | AF500075 | GCA_003605145.1 |
| *Shewanella upenei* | 20-23R$^{T}$ | GQ260190 | GCA_002836995.1 |
| *Tepidiphilus succinatimandens* | DSM 15512$^{T}$ | AY219713 | GCA_006503695.1 |
| *Tepidiphilus thermophilus* | JCM 19170$^{T}$ | HM543264 | GCA_001418245.1 |
| *Thalassospira permensis** | NBRC 106175$^{T}$ | FJ860275 | GCA_000714555.1 |
| *Thalassospira xiamenensis* | M-5$^{T}$ | CP004388 | GCA_000300235.2 |
| *Thermoanaerobacter brockii* subsp. *finnii* | Ako-1$^{T}$ | CP002466 | GCA_000175295.2 |
| *Thermoanaerobacter ethanolicus* | JW 200$^{T}$ | AEYS01000048 | GCA_003722315.1 |
| *Thermoanaerobacter italicus* | Ab9$^{T}$ | CP001936 | GCA_000025645.1 |
| *Thermoanaerobacter mathranii* subsp. *mathranii* | A3$^{T}$ | CP002032 | GCA_000092965.1 |
| *Thermoanaerobacter pseudethanolicus* | ATCC 33223$^{T}$ | CP000924 | GCA_000019085.1 |
| *Thermoanaerobacter siderophilus* | SR4$^{T}$ | CM001486 | GCA_000262445.1 |
| *Thermoanaerobacter thermocopriae* | JCM 7501$^{T}$ | L09167 | GCA_000518565.1 |
| *Thermoanaerobacter wiegelii* | Rt8.B1$^{T}$ | CP002991 | GCA_000147695.3 |
| *Thermodesulfovibrio islandicus* | DSM 12570$^{T}$ | AXWU01000024 | GCA_000482825.1 |
| *Thermodesulfovibrio yellowstonii* | DSM 11347$^{T}$ | CP001147 | GCA_000020985.1 |
| *Thermotoga naphthophila* | RKU-10$^{T}$ | ACXW01000001 | GCA_000025105.1 |
| *Thermotoga petrophila* | RKU-1$^{T}$ | CP000702 | GCA_000016785.1 |

*: *Corynebacterium ihumii* GD7$^{T}$, and *Thalassospira permensis* NBRC 106175$^{T}$ are not validly published species.

†: The whole genome sequence of *Rufibacter quisquiliarum* DSM 29854$^{T}$ was not available on NCBI and was downloaded from IMG (https://img.jgi.doe.gov). The number indicates the IMG taxon ID.

**Table S2.** Differential characteristics of *Actinokineospora mzabensis* PAL84$^T$ and *Actinokineospora spheciospongiae* EG49$^T$. Data from Lei *et al.* (2020). +, Positive; w, weakly positive; −, negative; nd, no data available.

| **Characteristic** | ***Actinokineospora mzabensis* PAL84$^T$** | ***Actinokineospora spheciospongiae* EG49$^T$** |
|---|---|---|
| Source | Soil | Sponge |
| Motility | – | – |
| Growth on ISP 3 | Good | Good |
| Aerial mycelium | Pinkish purple | White |
| Substrate mycelium | Purple to blackish | Yellow to tan |
| Use of sole carbon sources (1.0%): | | |
| Cellobiose | + | + |
| Galactose | + | – |
| Sucrose | – | + |
| Xylose | + | + |
| Rhamnose | + | w |
| Fructose | + | + |
| Maltose | + | + |
| Mannitol | + | + |
| Arabinose | – | – |
| Nitrate reduction | + | + |
| Gelatin liquefaction | + | + |
| DNA G+C content (mol%) | 72.8 | 72.8 |
| Major polar lipids* | PE | DPG, PE, OH-PE |
| Major fatty acids | iso-$C_{16:0}$, iso-$C_{15:0}$, iso-$C_{16:1}$ h, iso-$C_{16:0}$2OH | iso-$C_{16:0}$, iso-$C_{14:0}$, iso-$C_{15:0}$, iso-$C_{16:1}$ h |
| DNA G+C content (mol%) | 72.8 | 72.8 |
| Genome size (bp) | 7,546,603 bp | 7,529,476 bp |
| GenBank Accession | GCA_003182415.1 | GCA_000564855.1 |

*DPG, diphosphatidylglycerol; PG, phosphatidylglycerol; PI, phosphatidylinositol; PL, phospholipid; UL, unidentified lipid.

**Table S3.** Physiological and chemotaxonomic characteristics that differentiate *C. afermentans* subsp. *afermentans* DSM 44280$^T$ and *Corynebacterium ihumii* GD7$^T$. +, Positive; −, negative; nd, not determined; DPG, diphosphatidylglycerol; PE, phosphatidylethanolamine; PG, phosphatidylglycerol; PIM, phosphatidylinositol mannosides; PI, phosphatidylinositol; PL, unidentified phospholipid; GL, unidentified glycolipid; UL, unidentified lipid. Data from Atasayar et al. (2017). All strains were negative for: utilization of mannitol, lactose, sucrose, glycogen, production of β-galactosidase and N-acetyl-*β*-glucosaminidase, hydrolysis of aesculin and gelatin. All strains were positive for catalase. All strains contained corynemycolates.

| **Characteristic** | ***C. afermentans* subsp. *afermentans* DSM 44280$^T$** | ***Corynebacterium ihumii* GD7$^T$** |
|---|---|---|
| Utilization of: | | |
| Glucose | − | + |
| Ribose | − | − |
| Maltose | − | − |
| Xylose | − | + |
| Reduction of nitrate | − | − |
| Production of: | | |
| Pyrazinamidase | + | + |
| Pyrolidonyl arylamidase | + | − |
| Alkaline phosphatase | + | + |
| β-Glucuronidase | + | − |
| α-Glucosidase | − | − |
| Hydrolysis of: | | |
| Urea | − | − |
| Activity of | | |
| Oxidase | nd | − |
| Tuberculostearic acid | No | − |
| Corynemycolates | + | nd |
| Dominant fatty acids (>10% of total) | $C_{16:0}$, $C_{15:0}$, $C_{18:1}$ $\omega 9c$ | nd |
| Polar lipids | DPG, PG, PI, PIM, 3×GL, 6×UL | nd |
| Dominant menaquinones (>10% of total) | MK-8($H_2$), MK-9($H_2$), MK-(10 $H_2$) | nd |
| DNA G+C content (mol%) | 64.9 | 64.5 |
| Genome size (bp) | 2,326,687 bp | 2,251,282 bp |
| GenBank Accession | GCA_900156035.1 | GCA_000403725.1 |

**Table S4.** Phenotypic properties that distinguish strain *Dietzia papillomatosis* N 1280$^T$, *D. cinnamea* DSM 44904$^T$ and *D. maris* DSM 43672$^T$. All strains hydrolysed allantoin, arbutin and urea, were catalase-positive, reduced nitrate, were resistant to lysozyme, degraded cellulose, starch, Tweens 40 and 60, and grew at 37 °C and pH 7 and 10. Adonitol, l-arabinose, arbutin, butane-1,3-diol, butane-1,4-diol, butane-1-ol, butane-2,3-diol, d-cellobiose, dextrin, ethanol, meso-erythritol, d-fructose, d- and l-fucose, d-galactose, d-gentiobiose, d-glucose, glycerol, myo-inositol, d-lactose, maltose, d-mannitol, d-mannose, d-melezitose, d-melibiose, propane-1,2-diol, propane-1,3-diol, d-raffinose, d-ribose, d-salicin, d-sorbitol, sucrose, d-tagatose, trehalose, turanose, d-xylose and d-xylitol were used as sole carbon sources by all strains [all at 1.0% (w/v) or 1.0% (v/v)]. Acetamide, l-alanine, l-asparagine, l-aspartic acid, l-glutamic acid, l-glycine, l-histidine, l-leucine, l-isoleucine, dl-norleucine, l-norvaline, l-ornithine, dl-phenylalanine, l-proline, l-serine, l-thymidine, l-valine and urea were used as sole carbon and nitrogen sources. All strains were negative for aesculin hydrolysis, oxidase activity and degradation of casein, DNA, gelatin, hypoxanthine, pectin, RNA, xanthine and xylan. None of the strains utilized sodium adipate, sodium gluconate, sodium malonate, sodium oleate, sodium oxalate, sodium suberate or sodium succinate as sole carbon sources (all at 0.1%, w/v) or grew at pH 5.0. +, Positive; −, negative; w, weakly positive. Data from Jones et al. (2008).

| **Characteristic** | ***Dietzia papillomatosis* N 1280$^T$** | ***D. cinnamea* DSM 44904$^T$** | ***D. maris* DSM 43672$^T$** |
|---|---|---|---|
| Colony colour | Orange | Orange | Orange |
| Nitrite reduction | − | − | − |
| Degradation of: | | | |
| Chitin | + | − | + |
| Elastin | − | − | + |
| l-Tyrosine | + | − | + |
| Tributyrin | − | − | + |
| Uric acid | − | − | − |
| Acid production (aerobically) from: | | | |
| d-Fructose | + | + | + |
| d-Glucose | − | + | − |
| d-Mannose | − | + | − |
| d-Raffinose | − | − | w |
| Sucrose | − | + | + |
| Utilization as sole carbon and nitrogen sources: | | | |
| l-Arginine | − | + | − |
| l-Cysteine | + | + | + |
| Growth at: | | | |
| 5 °C | − | − | − |
| 10 °C | − | + | + |
| 45 °C | − | − | + |
| Growth in the presence of: | | | |
| 7% (w/v) NaCl | + | + | + |
| 8% (w/v) NaCl | + | + | − |
| DNA G+C content (mol%) | 70.9 | 70.8 | 70.9 |
| Genome size (bp) | 3,486,774 bp | 3,598,827 bp | 3,505,372 bp |
| GenBank Accession | GCA_001570845.1 | GCA_001571065.1 | GCA_001630765.1 |

**Table S5.** Physiological characteristics of *N. exalbida* IFM 0803$^T$ *and N. gamkensis* DSM 44956$^T$.

| ***Biochemical tests:*** | ***N. exalbida* IFM 0803$^T$** | ***N. gamkensis* DSM 44956$^T$** |
|---|---|---|
| Allantoin | + | nr |
| Arbutin | + | + |
| *Degradation tests:* | | |
| Esculin | + | + |
| Adenine | − | − |
| Arbutin | − | w |
| Casein | + | + |
| Elastin | − | nr |
| Hypoxanthine | − | − |
| Testosterone | + | nr |
| Tyrosine | − | + |
| Uric acid | + | nr |
| Xanthine | − | − |
| **Growth on sole carbon sources (1%, w/v):** | | |
| Adonitol | nr | − |
| L -(+)-Arabinose | − | − |
| D -(+)-Cellobiose | nr | w |
| *meso*-Erythritol | − | − |
| D -(–)-Fructose | nr | + |
| D -(+)-Galactose | − | w |
| Gluconate | − | nr |
| D -(+)-Glucose | + | + |
| Inositol | − | − |
| Inulin | nr | w |
| Lactose | nr | w |
| D -(+)-Maltose | − | nr |
| Mannitol | nr | − |
| D -(+)-Mannose | − | w |
| D -(+)-Melibiose | nr | w |
| D -(+)-Melezitose | nr | − |
| D -(+)-Raffinose | nr | − |
| L -(+)-Rhamnose | − | − |
| Sorbitol | − | nr |
| D -(+)-Sucrose | nr | − |
| D -(+)-Trehalose | nr | w |
| Xylitol | nr | w |
| D -(+)-Xylose | nr | − |
| **Growth on sole carbon sources (1%, w/v):** | | |
| Acetamide | − | nr |
| Sodium acetate | nr | + |
| Sodium adipate | − | nr |
| Sodium citrate | + | + |
| **Growth on sole nitrogen sources (0.1%, w/v):** | | |
| *l* -Histidine | nr | + |
| *l* -Phenylalanine | nr | + |
| *l* -Proline | nr | − |
| *l* -Serine | nr | + |
| *l* -Valine | nr | + |
| Cell wall | meso-A2pm; Ara, Gal (chemotype IV) | Cell wall: meso-A2pm; Ara, Gal (chemotype IV) |
| Polar lipid | nd | PE (type PII) |
| Quinone | MK-8(H4, ω-cycl) | nd |
| Mycolic acid | Mycolic acid (C48-56) | nd |
| DNA G+C content (mol%) | 68.6 | 68.4 |
| Genome size (bp) | 7,367,991 bp | 7,709,792 bp |
| GenBank Accession | GCA_000308575.1 | GCA_001612985.1 |

Data from Goodfellow *et al.* (2012). Symbols: +, positive reaction; −, negative reaction; w, weak reaction; nr, not reported.

**Table S6.** Physiological characteristics of *Nocardia coubleae* OFN N12$^{T}$ and *N. ignorata* DSM 44496$^{T}$. Data are taken from Isik et al. (1999); Yang et al. (2018); Rodríguez-Nava et al. (2007). −, Negative; +, positive; w, weakly positive; nd, not determined.

| **Characteristic** | ***Nocardia coubleae*** **OFN N12$^{T}$** | ***N. ignorata*** **DSM 44496$^{T}$** |
|---|---|---|
| Colony color | Orange to white | Orange |
| Temperature range (°C) | 25–37 | 22–45 |
| NaCl range (%, w/v) | 0–6 | 0–6 |
| pH range | 6–10 | 5–10 |
| Production of H2S | – | – |
| Reduction of nitrate | – | – |
| Starch | – | – |
| Gelatin | – | + |
| Urea | nd | + |
| Tween 40 | + | – |
| Tween 80 | + | + |
| Growth on carbon sources (%, w/v): | | |
| D-Glucose (1.0) | + | + |
| l-Arabinose (1.0) | − | − |
| d-Fructose (1.0) | + | + |
| d-Galactose (1.0) | + | w |
| Glycerol (1.0) | + | − |
| Maltose (1.0) | + | + |
| d-Mannitol (1.0) | + | w |
| Mannose (1.0) | + | + |
| Raffinose (1.0) | + | − |
| l-Rhamnose (1.0) | − | − |
| d-Ribose (1.0) | + | + |
| Sucrose (1.0) | + | − |
| Sorbitol (1.0) | − | − |
| d-Glucose | + | + |
| Lactose | + | – |
| *myo*-Inositol | – | – |
| Growth on Bennett's agar at: | | |
| 25 °C | | |
| 37 °C | + | + |
| 45 °C | − | − |
| Decomposition of (%, w/v): | | |
| Adenine (0.4) | − | − |
| Casein (1.0) | − | − |
| Hypoxanthine (0.4) | − | − |
| Testosterone (0.1) | + | + |
| Tyrosine (0.5) | − | − |
| Uric acid (0.5) | − | − |
| Cell wall | meso-A2pm; Ara, Gal, Glc, Rib | meso-A2pm; Ara, Gal (chemotype IV) |
| Quinone | MK-8(H4, ω-cycl) | MK-8(H6), 2,3-epoxy-MK-8(H6) |
| Polar lipid | nd | PE (type PII) |
| Mycolic acid | Mycolic acid (C52-58) | Mycolic acid (C46-54) |
| DNA G+C content (mol%) | 67.9 | 67.7 |
| Genome size (bp) | 6,619,824 bp | 7,022,989 bp |
| GenBank Accession | GCA_001612805.1 | GCA_004362495.1 |

**Table S7.** Differential physiological characteristics of strains *Nocardia elegans* IMMIB N-402$^T$ and *N. nova* ATCC 33726$^T$. All strains are positive for utilization of glucose as a carbon source and for hydrolysis of urea. All strains are negative for the following: hydrolysis of casein, elastin, gelatin, tyrosine, xanthine; utilization of cellobiose, raffinose, sorbitol, sucrose, meso-erythritol and myo-inositol as carbon sources; and utilization of acetamide, gelatin, proline and serine as simultaneous carbon and nitrogen sources. w, weakly utilized after 3 weeks incubation. Data from Yassin and Brenner (2005).

| **Characteristic** | ***Nocardia elegans* IMMIB N-402$^T$** | ***N. nova* ATCC 33726$^T$** |
|---|---|---|
| Hydrolysis of: | | |
| Aesculin | + | − |
| Hypoxanthine | − | − |
| Testosterone | − | − |
| Utilization as sole sources of carbon and energy | | |
| Acetate | + | + |
| Citrate | − | − |
| Gluconate | − | − |
| l-Arabinose | − | − |
| Galactose | − | + |
| Maltose | − | − |
| l-Rhamnose | − | − |
| Trehalose | − | − |
| Xylose | − | − |
| Mannitol | − | − |
| iso-Amyl alcohol | − | + |
| 2,3-Butanediol | − | − |
| 1,2-Propanediol | − | − |
| *m*-Hydroxybenzoate | − | − |
| *p*-Hydroxybenzoate | − | − |
| Utilization of l-alanine as sole source of carbon and nitrogen | − | − |
| Quinone | MK-8(H6, ω-cycl) | nd |
| Cell wall | meso-A2pm; Ara, Gal (type IV) [6086]. | nd |
| Polar lipid | PE, PI, PIM, DPG (type PII) | nd |
| DNA G+C content (mol%) | 67.9 | 67.9 |
| Genome size (bp) | 7,539,150 bp | 7,849,771 bp |
| GenBank Accession | GCA_001612845.1 | GCA_001613005.1 |

**Table S8.** Distinguishing characteristics of *Rhodococcus imtechensis* RKJ300$^T$ and *R. opacus* MTCC 6420$^T$. All of the strains were found to produce acid from glucose, fructose, d-galactose, inositol, lactose, maltose, d-mannitol, d-raffinose, sorbitol and sucrose and not to produce acid from adonitol, cellobiose or dulcitol. All strains were positive for the utilization of fructose, α-d-glucose, pyruvic acid and Tween 40 as sole sources of carbon and energy (Biolog). All strains were negative for the utilization of N-acetyl-l-glutamic acid, adenosine 5′-monophosphate, 2′-deoxyadenosine, β-dextrin, d-fructose 6-phosphate, α-d-glucose 1-phosphate, l-glutamic acid, glycogen, inosine, lactamide, mannan, methyl α-d-galactoside, thymidine, thymidine 5′-monophosphate and uridine as sole sources of carbon and energy (Biolog). Characteristics are scored as follows: +, positive; w, weakly positive; −, negative. Data from Ghosh et al. (2006).

| **Characteristic** | ***R. imtechensis* RKJ300$^T$** | ***R. opacus* MTCC 6420$^T$** |
|---|---|---|
| Hydrolysis of Tween 80 | + | − |
| Utilization of substrate as sole source of carbon and energy | | |
| l-Arabinose | − | − |
| l-Arabitol | − | + |
| d-Cellobiose | − | + |
| d-Maltose | + | + |
| d-Melezitose | − | + |
| Lactose | + | + |
| d-Raffinose | − | + |
| l-Rhamnose | + | − |
| d-Ribose | − | w |
| 2,4-Dinitrophenol | + | − |
| *p*-Nitrophenol | + | − |
| Utilization of substrate as sole source of nitrogen | | |
| l-Proline | + | − |
| l-Serine | + | + |
| Utilization of substrate as carbon source (using Biolog GP2 plate) | | |
| Acetic acid | + | − |
| Amygdalin | − | − |
| 2,3-Butanediol | − | + |
| *α*-Cyclodextrin | + | − |
| l-Fucose | + | − |
| d-Galacturonic acid | + | − |
| Gentiobiose | − | − |
| d-Lactic acid methyl ester | + | − |
| Lactulose | + | − |
| l-Malic acid | − | − |
| d-Melibiose | − | w |
| Methyl *β*-d-galactoside | − | − |
| Methyl *α*-d-mannoside | + | − |
| Propionic acid | + | − |
| d-Psicose | + | w |
| Putrescine | − | + |
| l-Pyroglutamic acid | + | − |
| Salicin | − | − |
| Stachyose | − | − |
| d-Trehalose | + | − |
| Uridine 5′-monophosphate | + | − |
| d-Xylose | − | + |
| DNA G+C content (mol%) | 67.2 | 67.3 |
| Genome size (bp) | 8,231,340 bp | 8,534,314 bp |
| GenBank Accession | GCA_000260815.1 | GCA_001646735.1 |

**Table S9.** Phenotypic characteristics that distinguish *Hfx. alexandrines, Hfx. lucentense* and ***Hfx. volcanii***. Data derived from Xu et (2007), and Allen et al. (2008). +, Positive; −, negative; w, weakly positive; nd, not determined.

| **Characteristic** | ***Hfx. alexandrinus*** | ***Hfx. lucentense*** | ***Hfx. volcanii*** |
|---|---|---|---|
| Motility | − | + | − |
| Colony colour | Red | Pink | Red to orange |
| NaCl range (M) | 1.7–5.2 | 1.8–5.1 | 1.0-4.5 |
| NaCl optimum (M) | 4.3 | 4.3 | 17-2.5 |
| Minimum $Mg^{2+}$ (M) | 0.33 | nd | nd |
| pH range | 5.5–7.5 | 5–9 | nd |
| pH optimum | 7.2 | 7.5 | 7 |
| Temp. range (°C) | 20–55 | 10–40 | nd |
| Temperature optimum (°C) | 37 | 37 | 45 |
| Oxidase test | + | + | + |
| Anaerobic growth on nitrate | − | nd | − |
| Nitrate reduction | + | − | + |
| $H_2S$ formation from thiosulfate | + | + | + |
| Hydrolysis of: | | | |
| Starch | − | − | − |
| Casein | − | − | − |
| Gelatin | + | − | + |
| Tween 80 | + | nd | − |
| Acid production from: | | | |
| Mannose | − | nd | − |
| Arabinose | + | + | + |
| Galactose | − | − | + |
| Xylose | + | + | + |
| Sucrose | + | − | + |
| Resistance to: | | | |
| Rifampicin | + | nd | − |
| Bacitracin | − | nd | − |
| DNA G+C content (mol%) | 66.3 | 66.4 | 65.5 |
| Genome size (bp) | 3,649,262 bp | 3,619,064 bp | 4,012,900 bp |
| GenBank Accession | GCA_000336735.1 | GCA_000336795.1 | GCA_000025685.1 |

**Table S10.** Differential characteristics of *Methanobacterium arcticum* M2$^T$ and *M. veterum* MK4$^T$. Data were obtained in this study unless otherwise indicated. All strains grew with $H_2/CO_2$ and formed filaments. Data from Shcherbakova et al. (2011).

| **Characteristic** | ***M. arcticum* M2$^T$** | ***M. veterum* MK4$^T$** |
|---|---|---|
| Cell shape | Rods, cyst-like cells | Rods |
| Cell dimensions (μm) | | |
| Width | 0.45–0.5 | 0.40–0.45 |
| Length | 3.0–6.0 | 2.0–8.0 |
| Substrate(s) | Formate | Methylamine/$H_2$, methanol/$H_2$ |
| Stimulatory factor(s) | None | Acetate |
| Temperature for growth (°C) | | |
| Range | 15–45 | 10–45 |
| Optimum | 37 | 28 |
| pH for growth | | |
| Range | 5.5–8.5 | 5.2–9.4 |
| Optimum | 6.8–7.2 | 7.0–7.2 |
| NaCl concentration for growth (M) | | |
| Range | 0–0.3 | 0–0.3 |
| Optimum | 0.1 | 0.05 |
| DNA G+C content (mol%) | 33.2 | 33.2 |
| Genome size (bp) | 3,369,555 bp | 3,393,923 bp |
| GenBank Accession | GCA_000745485.1 | GCA_000746075.1 |

**Table S11.** Characteristics of *Methanosarcina soligelidi* SMA-21$^T$ and *Methanosarcina mazei* DSM 2053$^T$ (Mah, 1980; Maestrojuán et al., 1992). +, positive; −, negative; ArPG, archaeol phosphatidylglycerol; Hydroxy-ArPG, hydroxyarchaeol phosphatidylglycerol; ArPE, archaeol phosphatidylethanolamine; Hydroxy-ArPE, hydroxyarchaeol phosphatidylethanolamine; nd, not determined.

| **Characteristic** | ***M. soligelidi* SMA-21$^T$** | ***M. mazei* DSM 2053$^T$** |
|---|---|---|
| Cell shape | Irregular cocci | Irregular cocci |
| Cell dimension (µm) | 1.3–2.5 | 1.0–3.0 |
| Gram stain | − | − |
| Temperature range for growth (°C) | 0–54 | 23–50 |
| Optimum temperature (°C) | 28 | 30–40 |
| pH range | 4.8–9.9 | 6.1–8.0 |
| Optimum pH | 7.8 | 7 |
| Tolerance of NaCl (M) | 0.02–0.6 | 0.1–1.0 |
| Optimum NaCl for growth | 0.02 | 0.1–0.3 |
| Utilization of: | | |
| $H_2/CO_2$ | + | + |
| Methanol | + | + |
| Acetate | + | + |
| Dimethyl sulfide | − | − |
| Monomethylamine | − | − |
| Dimethylamine | − | + |
| Trimethylamine Membrane lipids | − | + |
| ArPG | + | + |
| Hydroxy-ArPG | + | + |
| ArPE | + | + |
| Hydroxy-ArPE | + | + |
| DNA G+C content (mol%) | 41.5 | 41.4 |
| Genome size (bp) | 4,064,496 bp | 4,142,816 bp |
| GenBank Accession | GCA_000744315.1 | GCA_000970205.1 |

**Table S12.** Phenotypic and biochemical characteristics of strain *R. quisquiliarum* CAI-18b$^T$ and *R. ruber* CCM 8646$^T$. Cells are Gram-stain-negative, short rods, pinkish-red colonies, occurring predominantly in pairs or in irregular clusters and are non-spore-forming. Positive for catalase, esterase lipase (C8), leucine arylamidase, and valine arylamidase. +, Present; −, absent; w, weak reaction; nt, not tested.

| **Characteristic** | ***R. quisquiliarum* CAI-18b$^T$** | ***R. ruber* CCM 8646$^T$** |
|---|---|---|
| Motility | + | nt |
| Temperature range (optimum) (°C) | 4–45 (20–37) | 10 to 37 |
| pH range (optimum) | 7–11 (8 to 10) | 7 to 11 |
| NaCl concentration for growth (%) | 0–2 | 0-1 |
| Assimilation of (API 50 CH): | | |
| d-Arabinose | − | w |
| d-Galactose | + | - |
| d-Fructose | − | + |
| d-Mannose | + | + |
| N-Acetylglucosamine | + | + |
| Aesculin | + | + |
| Cellobiose | + | - |
| Lactose | + | - |
| Melibiose | + | - |
| Sucrose | − | + |
| Inulin | − | nt |
| Raffinose | − | + |
| Glycogen | + | + |
| Gentiobiose | + | + |
| dextrin | nt | + |
| α-D-glucose | + | + |
| maltose | + | + |
| D-trehalose | + | + |
| Enzyme activity (API ZYM) | | |
| Lipase (C14) | − | - |
| Cystine arylamidase | w | - |
| α-Chymotrypsin | − | - |
| α-Galactosidase | + | w |
| β-Galactosidase | − | − |
| α-Glucosidase | + | + |
| Predominant fatty acids (>10%) | iso-$C_{15:0}$ and iso-$C_{17:1}$ I | SF4 (iso-$C_{17:1}$ I/anteiso-$C_{17:1}$ B), iso-$C_{15:0}$, $C_{17:1}$ $\omega$6c and SF3 ($C_{16:1}$ $\omega$7c/$C_{16:1}$ $\omega$6c) |
| Polar lipids | PE and an unknown APL | PE, an unknown AG and six unknown PL. |
| Major respiratory quinone | MK-7 | MK-7 |
| DNA G + C content (mol%) | 51.7 | 51.5 |
| Genome size (bp) | 5589356 bp | 5,502,314 bp |
| GenBank Accession | JGI Project Id: 1227356 | GCA_001647275.1 |

Data from Felföldi et al. (2016); Ký́rova et al. (2016).

**Table S13.** Discriminating characteristics of subspecies of *Caldanaerobacter subterraneus* subsp. *tengcongensis* and *Caldanaerobacter subterraneus* subsp. *yonseiensis*. Both subspecies produced l-alanine (determined in this study), acetate, $H_2$ and $CO_2$ as diagnostic fermentation products from glucose. +, Positive; −, negative; nd, not determined.

| **Characteristic** | ***Caldanaerobacter subterraneus* subspecies** | |
|---|---|---|
| | ***tengcongensis*†** | ***yonseiensis*‡** |
| Type strain | JCM 11007$^T$ | DSM 13777$^T$ |
| Source | Hot spring | Geothermal water |
| **Temperature for growth (°C):** | | |
| Range | 50–80 | 50–85 |
| Optimum | 75 | 75 |
| pH for growth: | | |
| Range | 5·5–9·0 | 4·5–9·0 |
| Optimum | 7·0–7·5 | 6·5 |
| **NaCl concentration for growth (%):** | | |
| Range | 0–2·5 | 0–4 |
| Optimum | 0·2 | 0 |
| Use of CO | + | + |
| **Diagnostic fermentation products from glucose:** | | |
| Ethanol | + | + |
| Lactate | − | + |
| DNA G + C content (mol%) | 37.6 | 37.5 |
| Genome size (bp) | 2,689,445 bp | 2,700,546 bp |
| GenBank Accession | GCA_000007085.1 | GCA_000473865.1 |

†Data from Xue et al. (2001); ‡Data from Kim et al. (2001).

**Table S14.** Properties of *Carboxydothermus hydrogenoformans* Z-2901$^T$ and *C. ferrireducens* JW/AS-Y7$^T$. Data from Yoneda et al. (2012). nr, Not reported.

| **Characteristic** | ***C. hydrogenoformans* Z-2901$^T$** | ***C. ferrireducens* JW/AS-Y7$^T$** |
|---|---|---|
| Sampling site pH | 5.5 | 6.0–8.3 |
| Motility | + | + |
| Growth temperature (°C) | | |
| Range | 40–78 | 50–74 |
| Optimum | 70–72 | 65 |
| Growth pH | | |
| Range | 6.6–8.0 | 5.5–7.6 |
| Optimum | 7 | 6.0–6.2 |
| DNA G+C content (mol%) | 39–41 | 41 |
| Major CFAs (% of total) | | |
| $C_{14:0}$ | 11.3 | 7.4 |
| iso-$C_{15:0}$ | 30.1 | 30.8 |
| $C_{15:0}$ | 2.3 | 1.9 |
| $C_{16:0}$ | 20.2 | 24.4 |
| Electron acceptor with CO as electron donor† | | |
| Ferric citrate | + (−) | − (+‡) |
| Amorphous iron (III) oxide | − (−) | − (+‡) |
| AQDS | −(+) | − (+†) |
| Sulfate | − (−) | − (−) |
| Sulfite | − | − |
| Thiosulfate | − | − |
| Elemental sulfur | − (−) | − |
| Nitrate | − | − |
| Fumarate | − (+) | − (+‡) |
| None | − (+) | − (−) |
| DNA G + C content (mol%) | 42 | 41.9 |
| Genome size (bp) | 2,401,520 bp | 2,441,992 bp |
| GenBank Accession | GCA_000012865.1 | GCA_000427565.1 |

†Data are from this study apart from data in parentheses, which are taken from previous reports.

‡Carboxydotrophic growth without production of $H_2$.

**Table S15.** Differential phenotypic characteristics between *Carnobacterium inhibens* subsp. *gilichinskyi* and *C. inhibens* K1$^T$. Data from Nicholson et al. (2015). +, Positive; −, negative; (+) weakly positive; nd, not determined.

| **Characteristic** | ***C. inhibens* subsp. *gilichinskyi*** | ***C. inhibens* K1$^T$** |
|---|---|---|
| Isolation source | Siberian permafrost | Atlantic salmon |
| Temperature range (optimum) (°C) | 0–40 (25) | 0–30 |
| pH range for growth (optimum) | 5.5–9.0 (8.2) | 5.5–9.0 |
| NaCl range for growth (optimum) (%) | 0–8.8 (0.5) | 0–6 |
| Growth on: | | |
| Aesculin | nd | + |
| d-Arabinose | nd | nd |
| l-Arabinose | nd | − |
| Erythritol | nd | − |
| d-Galactose | − | + |
| Gentiobiose | − | + |
| d-Glucose | − | + |
| Inositol | − | − |
| Lactose | − | (+) |
| Lithium chloride | − | + |
| Maltose | + | + |
| d-Mannitol | − | + |
| d-Mannose | + | + |
| Pectin | − | + |
| Potassium tellurite | + | − |
| Starch | + | − |
| Sucrose | + | + |
| Tetrazolium Blue | − | + |
| Trehalose | + | + |
| DNA G+C content (mol%) | 35.2 | 34.9 |
| Genome size (bp) | 2,497,115 bp | 2,748,608 bp |
| GenBank Accession | GCA_000493735.1 | GCA_000746825.1 |

**Table S16.** Characteristics differentiating *Geobacillus kaustophilus* and *Geobacillus thermoleovorans*. +, positive; −, positive; w, weak reaction; v, result varies within strains; nd, no data available. All strains were negative for indole production. All strains were positive for catalase and production of acid from glucose, fructose, maltose and mannose. Data from Semenova et al. (2019).

| Characteristic | *Geobacillus kaustophilus* | *Geobacillus thermoleovorans* |
|---|---|---|
| Cell length (µm) | ≤3 | 2.0–6.0 |
| Cell width (µm) | ≤0.9 | 0.7–1.5 |
| Motility | nd | v |
| Spores: | | |
| Cylindrical | − | − |
| Subterminal | + | + |
| Terminal | + | + |
| Central/paracentral | − | − |
| Acid production from: | | |
| *N*-Acetylglucosamine | nd | v |
| Amygdalin | nd | − |
| Adonitol | + | − |
| l-Arabinose | − | − |
| Arbutin | nd | − |
| Cellobiose | + | v |
| Galactose | v | v |
| Gentibiose | nd | − |
| Glycerol | − | + |
| Glycogen | nd | v |
| Inositol | + | v (myo) |
| Lactose | − | v (w) |
| Mannitol | + | v |
| Melibiose | nd | v |
| Methyl d-glucoside | nd | v |
| d-Raffinose | nd | v |
| Rhamnose | − | − |
| d-Ribose | − | + |
| Salicin | nd | w |
| Sorbitol | − | − |
| Sucrose | + | v |
| Trehalose | + | v |

| | | |
|---|---|---|
| Turanose | nd | − |
| d-Xylose | + | v |
| Hydrolysis of: | | |
| Aesculin | − | + |
| Casein | + | v |
| Gelatin | + | + |
| ONPG | nd | v |
| Starch | + | + |
| Utilization of: | | |
| *n*-Alkanes ($C_{10}$–$C_{16}$) | nd | nd |
| Formate | v | nd |
| Acetate | − | nd |
| Lactate | − | + |
| Citrate Simmons | + | − |
| Ethanol | nd | nd |
| Anaerobic growth | nd | w |
| Gas from glucose | − | nd |
| $NO_3^- \rightarrow NO_2^-$ | + | + |
| Gas from nitrate | − | nd |
| Methyl red test | + | nd |
| Voges–Proskauer reaction | − | v (w) |
| Oxidase activity | nd | + |
| Urease | − | − |
| Temperature range for growth (°C) | 37–68 | 37–70 |
| NaCl range for growth (%, w/v) | 0–5 | ≤0.5–1 |
| pH range for growth | 6.0–8.0 | 5.0–9.0 |
| DNA G+C content (mol%) | 51.9 | 52.3 |
| Genome size (bp) | 3,670,957 bp | 3,499,317 bp |
| GenBank Accession | GCA_005160065.1 | GCA_001610955.1 |

**Table S17.** Differentiating phenotypic characteristics of strain *Geobacillus lituanicus* N-3$^T$ and *G. stearothermophilus*. +, All strains are positive; v, characteristic is variable; −, all strains are negative; nd, not determined. Data were obtained from Kuisiene et al. (2004). All strains were positive for utilization of glucose, fructose, maltose, mannose and sucrose.

| **Characteristic** | ***G. lituanicus* N-3$^T$** | ***G. stearothermophilus*** |
|---|---|---|
| Temperature range (°C) | 55–70 | 37–65 |
| pH | 6·5 | ND |
| Motility | + | + |
| Catalase | + | V |
| Oxidase | + | V |
| Hydrolysis of: | | |
| Casein | + | V |
| Collagen* | + | + |
| Starch | + | + |
| Gas from nitrate | − | V |
| Denitrification | + | V |
| NaCl range (%) | 0–0·5 | 0–5·0 |
| Resistance to lysozyme | − | − |
| Production of acid from: | | |
| Arabinose | + | V |
| Cellobiose | + | − |
| Galactose | + | − |
| Mannitol | + | V |
| Ribose | + | ND |
| Xylose | + | V |
| DNA G+C content (mol%) | 52.1 | 53.1 |
| Genome size (bp) | 3,499,511 bp | 2,630,829 bp |
| GenBank Accession | GCA_002243605.1 | GCA_001277805.1 |

**Table S18.** Phenotypic characteristics that differentiate *Geobacillus yumthangensis* AYN2$^T$, *Geobacillus galactosidasius* CF1B$^T$ and *Parageobacillus toebii* DSM 14590$^T$. +, Positive; −, negative; nd, not determined. Data from Poli *et al.* (2011); Najar et al. (2018).

| **Characteristic** | ***G. yumthangensis* AYN2$^T$** | ***G. galactosidasius* CF1B$^T$** | ***P. toebii* DSM 14590$^T$** |
|---|---|---|---|
| Cell width (μm) | 0.4–0.6 | nd | 0.5–0.9 |
| Cell length (μm) | 2.5–5 | nd | 2–3.5 |
| Motility | + | nd | + |
| **Production of acid from:** | | | |
| Adonitol | nd | nd | − |
| l-Arabinose | − | nd | − |
| Cellobiose | + | + | − |
| Galactose | + | + | − |
| Ribose | + | − | − |
| Glycerol | + | − | − |
| Inositol | + | nd | + |
| Lactose | + | + | − |
| Rhamnose | + | nd | − |
| Sorbitol | + | nd | − |
| d-Xylose | − | + | − |
| **Hydrolysis of:** | | | |
| Gelatin | + | − | − |
| Casein | − | − | + |
| Starch | + | − | − |
| Aesculin | − | nd | − |
| **Utilization of:** | | | |
| Glucose | | − | − |
| Formate | + | nd | − |
| Acetate | + | + | − |
| Lactate | + | nd | − |
| Citrate | − | − | − |
| Fermentation of glucose | + | nd | − |
| Methyl red test | − | nd | − |
| Denitrification | nd | + | + |
| NaCl concentration for growth (%, w/v) | 0–5 | 0.1–0.2 | 0–5 |
| pH range for growth | 6.0–10 | 7.2 | 6.0–9.0 |
| Temperature range for growth (°C) | 40–70 | 50–75 | 45–70 |
| Oxidase | nd | + | − |
| **Fatty acids** | | | |
| *iso*C 15:0 | 12.8 | 36 | 33.1 |
| *iso*C 16:0 | 13.9 | 11.3 | 20.5 |
| *iso*C 17:0 | 13.8 | 37.7 | 33.4 |
| DNA G+C content (mol%) | 42.1 | 41.6 | 42.4 |
| Genome size (bp) | 3,409,966 bp | 3,794,830 bp | 3,323,060 bp |
| GenBank Accession | GCA_002494375.1 | GCA_002217735.1 | GCA_003688615.1 |

**Table S19.** Major characteristics that differentiate *Megamonas funiformis* YIT 11815$^T$ and *Megamonas rupellensis* FM1025$^T$. Data from Sakon et al (2008) and Chevrot et al. (2008). Both organisms are positive for acid production from glucose, maltose, lactose, mannose, arabinose, xylose and raffinose. +, Positive; −, negative; w, weakly positive.

| **Characteristic** | ***Megamonas funiformis* YIT 11815$^T$** | ***Megamonas rupellensis* FM1025$^T$** |
|---|---|---|
| Isolation source | Human faeces | caecum of a duck |
| Morphology | Very large rods | Very large rods |
| Cell size (µm) | 1×5–200 | 1.0–6.0 µm |
| Acid production from: | | |
| Melibiose | − | nd |
| Amygdalin | − | nd |
| Cellobiose | − | + |
| Inositol | − | nd |
| Glycerol | − | + |
| Melezitose | − | − |
| Rhamnose | − | − |
| Salicin | + | + |
| Hydrolysis of aesculin | − | nd |
| *α*-Mannosidase | − | nd |
| Haemolysis | Weak *β* (horse blood) | nd |
| DNA G+C content (mol%) | 31.5 | 31.3 |
| Genome size (bp) | 2,568,766 bp | 2,330,046 bp |
| GenBank Accession | GCA_010669225.1 | GCA_000378365.1 |

**Table S20.** Differential phenotypic characteristics of *Salimicrobium jeotgali* MJ3$^T$ and *S. salexigens* 29CMI$^T$ (data from de la Haba et al., 2011). All strains are positive for the following characteristics: hydrolysis of starch, utilization of d-glucose, d-mannose and sucrose, oxidase, catalase, alkaline phosphatase, esterase (C4), esterase lipase (C8), lipase (C14), leucine arylamidase, valine arylamidase, α-chymotrypsin, acid phosphatase, susceptibility to oleandomycin and acid production from l-arabinose, d-galactose, d-glucose and d-mannose. All strains are negative for the following characteristics: hydrolysis of Tween 80, casein and tyrosine, α-galactosidase, N-acetyl-β-glucosaminidase, α-fucosidase, α-mannosidase, assimilation of N-acetylglucosamine, caprate, adipate, malate, citrate and phenylacetate, susceptibility to novobiocin, lincomycin, gentamicin, neomycin and tetracycline, and acid production from d-mannitol, d-sorbitol and sucrose. +, Positive; −, negative; w, weakly positive.

| **Characteristic** | ***Salimicrobium jeotgali* MJ3$^T$** | ***S. salexigens* 29CMI$^T$** |
|---|---|---|
| Morphology | Cocci | Cocci |
| Motility | + | − |
| Gram staining | + | + |
| Spore formation | − | − |
| Colony colour | Yellow | Yellow |
| Nitrate reduction | + | + |
| Optimal NaCl concentration (%) | 10 | 7.5–12.5 |
| Hydrolysis of Tween 20* | + | + |
| Enzyme activities (API ZYM)* | | |
| Cystine arylamidase | w | w |
| Trypsin | − | w |
| Naphthol-AS-BI-phosphohydrolase | − | − |
| β-Galactosidase | − | − |
| α-Glucosidase | + | + |
| Susceptibility to:* | | |
| Kanamycin | − | − |
| Ampicillin | − | − |
| Carbenicillin | + | + |
| Utilization of:* | | |
| d-Xylose | + | − |
| Lactose | w | − |
| d-Fructose | + | − |
| l-Arabinose | − | − |
| d-Galactose | + | + |
| d-Mannitol | + | + |
| d-Sorbitol | + | − |
| Acid production from:* | | |
| d-Xylose | − | − |
| Lactose | + | − |
| d-Fructose | − | − |
| DNA G+C content (mol%) | 46.3 | 46.7 |
| Genome size (bp) | 2,776,927 bp | 2,584,100 bp |
| GenBank Accession | GCA_001685435.3 | GCA_900156705.1 |

**Table S21.** Distinguishing properties among *Thermoanaerobacter* species. Data from Xue et al. (2001). +, Positive; −, negative; nr, not reported; +w, weakly positive.

| **Feature** | ***T. ethanolicus*** | ***T. wiegelii*** | ***T. italicus*** | ***T. thermocopriae*** | ***T. siderophilus*** | ***T. mathranii.*** | ***T. pseudoethanolicus*** | ***T. brockii subsp. finnii* DSM 3389$^T$** |
|---|---|---|---|---|---|---|---|---|
| Gram reaction | v | – | – | – | + | v | v | v |
| Spores | – | + | + | + | + | + | + | + |
| Cell wall peptidoglycan diamino acid | *m*-DAP | nr | *m*-DAP | nr | nr | nr | nr | *m*-DAP |
| Motility | + | + | – | + | + | + | + | + |
| Optimum temperature ( C) | 69 | 66 | 70 | 60 | 70 | 70 | 65 | 65 |
| DNA G+C content (mol%) | 32 | 35 | 34 | 37 | 32 | 37 | 34.4 | 32 |
| H2 inhibition | nr | nr | nr | nr | – | + | nr | nr |
| Starch hydrolysis | + | + | + | + | + | nr | nr | + |
| Xylan degradation | + | nr | + | + | – | + | nr | nr |
| Growth substrates: | | | | | | | | |
| Glucose | + | + | + | + | + | + | + | + |
| Sucrose | + | + | + | – | + | + | + | + |
| Ribose | + | – | nr | – | nr | + | nr | + |
| Xylose | + | + | + | – | + | + | + | + |
| Arabinose | – | – | + | nr | – | + | nr | - |
| Mannitol | – | + | nr | + | nr | + | nr | + |
| Glycerol | – | + | nr | – | + | – | nr | - |
| Pyruvate | + | – | nr | nr | + | nr | nr | + |
| Fermentation products: | | | | | | | | |
| Acetate | + | + | + | + | – | + | nr | + |
| Lactate | + | + | + | + | + | + | nr | + |
| Ethanol | + | + | + | + | + | + | + | + |
| Antibiotic inhibition: | | | | | | | | |
| Chloromycetin | + | – | nr | nr | + | + | nr | nr |
| Penicillin | – | – | nr | nr | – | + | nr | + |
| Streptomycin | – | – | nr | nr | – | nr | nr | nr |
| Tetracycline | – | + | nr | nr | nr | + | nr | + |
| DNA G+C content (mol%) | 34.2 | 34.3 | 34.1 | 34.1 | 34.2 | 34.3 | 34.5 | 34.5 |
| Genome size (bp) | 2,911,280 bp | 2,785,056 bp | 2,451,061 bp | 2,455,744 bp | 2,540,159 bp | 2,306,092 bp | 2,362,816 bp | 2,344,824 bp |
| GenBank Accession | GCA_003722315.1 | GCA_000147695.3 | GCA_000025645.1 | GCA_000518565.1 | GCA_000262445.1 | GCA_000092965.1 | GCA_000019085.1 | GCA_000175295.2 |

**Table S22.** Differential phenotypic characteristics between *Thermodesulfovibrio yellowstonii* YP87$^T$ and *Thermodesulfovibrio islandicus* R1Ha3$^T$. All strains give positive results for the utilization of pyruvate, hydrogen (plus acetate) and formate (plus acetate) as electron donors (and carbon source) in the presence of sulfate. All strains give negative results for the utilization of acetate and ethanol as electron donors. All strains are positive for fermentative growth with pyruvate, but negative for fermentative growth with lactate. All strains are positive for utilization of sulfate and thiosulfate as electron acceptors, but negative for elemental sulfur. Data from Sekiguchi et al. (2008). −, Negative; +, positive; i, iso; ai, anteiso; nd, no data available.

| **Characteristic** | ***T. yellowstonii* YP87$^T$** | ***T. islandicus* R1Ha3$^T$** |
|---|---|---|
| Motility | + | + |
| Cell width (μm) | 0.3 | 0.4 |
| Cell length (μm) | 1.5 | 1.7 |
| Growth temperature (°C) | | |
| Range | 40–70 | 45–70 |
| Optimum | 65 | 65 |
| pH for growth | | |
| Range | 5.5–8.5 | nd |
| Optimum | 6.8–7.0 | nd |
| Major cellular fatty acids | i-$C_{16:0}$, i-$C_{17:0}$, ai-$C_{15:0}$ | nd |
| Utilization of lactate: | | |
| With sulfate | + | + |
| In the presence of methanogens | + | nd |
| Utilization of external electron acceptors | | |
| Sulfite | + | − |
| Nitrate | − | + |
| Fe(III) NTA | + | nd |
| Isolation source | Hydrothermal vent water | Slightly alkaline thermal spring |
| DNA G+C content (mol%) | 34.1 | 34.2 |
| Genome size (bp) | 2,003,803 bp | 2,048,874 bp |
| GenBank Accession | GCA_000020985.1 | GCA_000482825.1 |

**Table S23.** Characteristics that differentiate the *A. salmonicida* subspecies. -+, >85% positive; -, <15% positive; v, variable (15–85%). Data from Pavan et al. (2000).

| **Test** | ***A. salmonicida* subsp. *masoucida*** | ***A. salmonicida* subsp. *salmonicida*** |
|---|---|---|
| Motility | + | + |
| DL-Lactate utilization | - | - |
| urocanic acid utilization | + | + |
| sucrose fermentation | + | + |
| LDC | v | v |
| D-sorbitol fermentation | v | v |
| brown diffusible pigment | - | + |
| Growth at 35-37 °C | - | - |
| indole | + | - |
| Voges-Proskauer at 25 °C | + | - |
| KCN | - | - |
| Mannitol fermentation | + | + |
| sucrose fermentation | + | - |
| Gas from glucose | + | - |
| DNA G+C content (mol%) | 58.8 | 58.3 |
| Genome size (bp) | 4,502,258 bp | 4,934,557 bp |
| GenBank Accession | GCA_000647955.1 | GCA_900445115.1 |

**Table S24.** Characteristics that differentiate *Alteromonas addita* and *A. stellipolaris*. All species/strains are Gram-negative, motile, oxidase- and catalase-positive and negative for indole and H2S production, grow at 3–6% NaCl and produce lipase (Tween 80). +, Positive; −, negative; w, weak reaction; v, variable; nd, no data. Data from Van Trappen et al. (2004); Ivanova et al. (2005).

| **Characteristic** | ***A. addita*** | ***A. stellipolaris*** |
|---|---|---|
| Production of pigments | − | + |
| Growth at: | | |
| 4 °C | + | + |
| 40 °C | − | − |
| Nitrate reduction to nitrite | − | − |
| Growth in NaCl at: | | |
| 10% | + | + |
| 15% | − | − |
| Hydrolysis of: | | |
| Gelatin | w | + |
| Agar | w | − |
| Starch | + | + |
| Haemolysis | + | − |
| Assimilation of: | | |
| d-Mannitol | − | + |
| l-Lactate | + | − |
| G+C content (mol%) | 43.6 | 43.5 |
| Genome size (bp) | 4,649,146 bp | 4,904,192 bp |
| GenBank Accession | GCA_001562195.1 | GCA_001562115.1 |

**Table S25.** Differential characteristics of the *B. bronchiseptica* (Vandamme et al., 1996 ; Weyant et al., 1996 ), *B. parapertussis* (Vandamme et al., 1996 ; Weyant et al., 1996 ) and *B. pertussis* (Vandamme et al., 1996 ; Weyant et al., 1996 ). +, Positive; –, negative; W, weakly positive; v, strain-dependent; nd, no data available. All taxa are negative for acid production from d-glucose and assimilation of d-glucose, lactose, sucrose, maltose and d-mannitol.

| **Characteristic** | ***B. bronchiseptica*** | ***B. parapertussis*** | ***B. pertussis*** |
|---|---|---|---|
| Isolation source | Mammalian respiratory tract | Human respiratory tract | Human respiratory tract |
| Motility | + | − | − |
| Oxidase | + | − | + |
| Growth on/at: | | | |
| MacConkey agar | + | + | − |
| Simmons' citrate agar | + | v | − |
| 25 °C | + | v | − |
| 42 °C | v | v | − |
| Reduction of nitrate to nitrite (API 20NE) | + | − | − |
| Urease activity | + | + | − |
| Brown pigment on HI agar with tyrosine | − | + | − |
| Litmus milk alkalization | + | − | nd |
| Assimilation of: | | | |
| d-Xylose | − | − | − |
| d-Gluconate | − | − | − |
| Caprate | v | − | − |
| Adipate | + | − | − |
| l-Malate | v | − | − |
| Phenylacetate | + | − | − |
| Enzyme activity | | | |
| Alkaline phosphatase | v | − | − |
| Esterase lipase (C8) | v | + | + |
| Lipase (C14) | − | − | − |
| Chymotrypsin | − | + | + |
| Valine arylamidase | − | − | − |
| Naphthol-AS-BI-phosphohydrolase | − | − | + |
| DNA G+C content (mol%) | 68.2 | 68.1 | 67.7 |
| Genome size (bp) | 5,173,845 bp | 4,800,737 bp | 4,043,846 bp |
| GenBank Accession | GCA_900445725.1 | GCA_001525545.2 | GCA_000306945.1 |

**Table S26.** The difference in phenotypic and biochemical characteristics between strain *C. manganoxidans* HS$^T$ and *C. taiwanensis* On1$^T$

| **Characteristic** | ***C. manganoxidans* HS$^T$** | ***C. taiwanensis* On1$^T$** |
|---|---|---|
| Optimum growth temperature (°C) | 50 | 55 |
| Optimum growth pH | 8–9 | 7 |
| Cell size (μm) | 0.5–0.7×2.2–3.5 | 0.6–0.8×1.2–2.2 |
| Oxidase | + | — |
| Catalase | + | w |
| Utilization of: | | |
| Galactose | + | — |
| Malate | + | — |
| Malonate | + | — |
| Sucrose | + | — |
| Acetate | — | + |
| Fructose | — | + |
| Trehalose | — | + |
| **Fatty acids:** | | |
| 10:0 3OH | 4.4 | 6.3 |
| 12:00 | 3.6 | 4.3 |
| 14:00 | 3 | 2.9 |
| 15:1 *ω*6c | 1.1 | |
| **16:00** | **27.3** | **30.4** |
| 17:0 cyclo | 19.4 | 1.9 |
| 17:00 | 4.1 | |
| **18:1 *ω*7c** | **20** | **20** |
| 18:00 | | 2.1 |
| **Summed feature 3b** | **12.3** | **31.3** |
| G+C content (mol%) | 66.2 | 65.9 |
| Genome size (bp) | | |
| GenBank Accession | | |

[a]The data of *C. manganoxidans* HS$^T$ was obtained from Takeda et al. [22].

[a]Values are shown as a percentage of the total fatty acid content for each strain. All strains grew at 55 °C for 48 h on TSA medium, and then the fatty acid composition was analyzed. Values for fatty acid present at level of less than 1.0% in the strain are not given.

[b]Summed feature 3 comprises 16:1ω7c or 15:0 iso 2OH or both.

**Table 27.** Biochemical tests for the differentiation of species and subspecies of the *Cronobacter dublinensis* subsp. *lactaridi* and *C. dublinensis* subsp. *lausannensis*. Data from Iversen et al. (2008). +, >90% Positive; v, 20–80% positive; −, <10% positive. All results are from Biotype 100 unless otherwise indicated.

| **Characteristic** | ***Cronobacter dublinensis* subsp. *lactaridi*** | ***C. dublinensis* subsp. *lausannensis*** |
|---|---|---|
| Indole production* | + | v |
| Carbon source utilization: | | |
| Dulcitol† | − | − |
| Lactulose | + | − |
| Malonate‡ | − | − |
| Maltitol | + | − |
| Palatinose | + | + |
| Putrescine | + | v |
| Melezitose | − | − |
| Turanose | v | − |
| *myo*-Inositol† | + | − |
| *cis*-Aconitate | + | + |
| *trans*-Aconitate | + | + |
| 1-0-Methyl *α*-d-glucopyranoside | + | + |
| 4-Aminobutyrate | + | + |
| GenBank Accession | GCA_000409345.1 | GCA_000409365.1 |
| Genome size (bp) | 4,453,586 bp | 4,613,339 bp |
| G+C content (mol%) | 58.30% | 57.9% |

**Table 28.** Comparison of physiological and cytological properties of *Desulfotignum phosphitoxidans* FiPS-3 and *D. balticum*, Data from Kuever et al. (2001); Schink et al. (2002).

| **Property** | ***Desulfotignum phosphitoxidans* FiPS-3$^T$** | ***D. balticum* strain Sax$^T$** |
|---|---|---|
| Oxidation of | | |
| Phosphite | + | – |
| Formate | + | + |
| $H_2/CO_2$ | + | + |
| Acetate | – | + |
| Fumarate | + | + |
| Maleate | w | + |
| Malate | + | + |
| Succinate | w | + |
| Lactate | – | + |
| Pyruvate | + | + |
| Butyrate | – | + |
| Crotonate | – | + |
| Glutarate | – | + |
| 2-Oxoglutarate | – | + |
| 3-Oxoglutarate | – | + |
| Glutamate | + | + |
| Glucose | + | – |
| Arabinose | + | – |
| Xylose | + | – |
| Glycine | + | + |
| Betaine | + | + |
| Proline | + | + |
| Choline | – | n.d. |
| Yeast extract | + | + |
| Reduction of | | |
| Sulfite | + | + |
| Thiosulfate | + | + |
| Sulfur | – | – |
| Nitrate | – | – |
| Desulfoviridin | – | – |
| G+C content (mol%) | 51.3 | 51.2 |
| Genome size (bp) | 4,998,761 bp | 5,118,755 bp |
| GenBank Accession assembly | GCA_000350545.1 | GCA_000421285.1 |

**Table 29.** Differential characteristics of the species of the genus *Haemophilus*. Data from Kilian (2015).

| **Characteristics** | ***H. influenzae*** | ***H. aegyptius*** |
|---|---|---|
| V-factor requirement | + | + |
| ALA→porphyrins | - | - |
| Indole productiona | d | - |
| Ureasea | d | + |
| Ornithine decarboxylasea | d | - |
| Arginine dihydrolase | - | - |
| Lysine decarboxylase | d | - |
| H2S (lead acetate) | - | - |
| Hemolysis | - | - |
| D-Glucose, acid production | + | w |
| D-Glucose, gas production | - | - |
| *Acid from:* | | |
| D-Fructose | - | - |
| Sucrose | - | - |
| Lactose | - | - |
| D-Xylose | + | - |
| D-Ribose | + | w |
| D-Mannose | - | - |
| D-Mannitol | - | - |
| D-Sorbitol | - | - |
| L-Arabinose | - | - |
| L-Rhamnose | - | - |
| D-Galactose | + | w |
| Sorbose | - | - |
| Cellobioase | - | - |
| Maltose | + | w |
| Melibiose | - | - |
| Trehalose | - | - |
| Melizitose | - | - |
| Raffinose | - | - |
| Inulin | - | - |
| Dulicitol | - | - |
| Glycerol | - | - |
| meso-Erythritol | - | - |
| inositol | - | - |
| Xylitol | - | - |
| Esculin, Salicin, adonitol | - | - |
| β-Galactosidase (ONPG test)b | - | - |
| α-Galactosidase | - | - |
| α-Glucosidase | - | - |
| β-Glucosidase | - | - |
| α-Mannosidase | - | - |
| β-Xylosidase | - | - |
| β-Glucoronidase | d | - |
| α-Fucosidase | - | - |
| Catalase | + | + |
| oxidase | + | + |
| $CO_2$ enhances growth | - | - |
| Alkaline phosphatase | + | + |
| IgA1 protease | + | + |
| Nitrate reduction | + | + |
| Nitrite reduction | - | - |

**Table_S30.** Differential characteristics of *Methylomicrobium agile* ATCC 35068 and *Methylomicrobium album* BG8. Data from Orata *et al.* (2018).

| **Characteristic** | ***Methylomicrobium agile* ATCC 35068** | ***Methylomicrobium album* BG8** |
|---|---|---|
| Pigmentation | W to SL | W to SL |
| Motility | + | + |
| Cyst formation | - | - |
| Dessiccation resistance | - | - |
| Growth occurs with 3% NaCl | - | - |
| Temp. growth, range (°C) | 10 to 37 | 10 to 37 |
| Temp. growth, optimum (°C) | 25 to 30 | 25 to 30 |
| Growth at 37 °C | + | V |
| Growth at 45 °C | - | - |
| Heat resistance (80 °C) | - | - |
| pH growth, range | 6 to 9 | 6 to 9 |
| pH growth, optimum | 7 | 7 |
| pMMO | + | + |
| sMMO | - | - |
| Main fatty acid | C16:1 ϖ5t | C16:1 ϖ5t |
| GeneBank accession | GCA_000733855.1 | GCA_000214275.3 |
| G+C content (mol%) | 56.2 | 56.2 |
| Genome size (bp) | 4,526,071 bp | 4,493,444 bp |

**Table 31** Characteristics that differentiate *P. agarilytica* comb. nov. KCTC 12755T (Yong et al., 2007; Park & Yoon, 2013) and *P. chathamensis* comb. nov. JCM 13645T (Matsuyama et al., 2006; Park & Yoon, 2013). −, Negative; +, positive; w, weakly positive; nd, no data available. All strains are non-pigmented and motile and hydrolyse Tween 80 and gelatin but not urea.

| Characteristic | *Paraglaciecola agarilytica* KCTC 12755T | *Paraglaciecola chathamensis* JCM 13645T |
|---|---|---|
| Growth at 4 °C | − | + |
| Temperature range for growth (°C) | 7–30 | 4–30 |
| NaCl tolerance (%, w/v) | 2–8 | 1–10 |
| Hydrolysis of: | | |
| Aesculin | + | + |
| Casein | + | + |
| DNA | + | nd |
| ONPG | − | + |
| Starch | + | + |
| Nitrate reduction | − | − |
| Utilization of carbon compounds | | |
| Acetate | + | − |
| Arabitol | nd | nd |
| l-Arabinose | nd | − |
| Cellobiose | + | − |
| d-Galactose | + | + |
| d-Glucose | − | nd |
| *myo*-Inositol | nd | + |
| dl-Lactate | nd | − |
| Maltose | − | + |
| d-Mannitol | + | + |
| d-Mannose | + | + |
| Raffinose | nd | + |
| l-Rhamnose | nd | + |
| Sucrose | − | + |
| Trehalose | nd | + |
| l-Alanine | nd | + |
| l-Leucine | nd | + |
| l-Proline | nd | + |
| l-Glutamate | nd | + |
| Succinate | nd | w |
| Propionate | nd | w |
| Major lipids* | PE, PG | PE, PG |

*DPG, Diphosphatidylglycerol; PA, phosphatidic acid; PE, phosphatidylethanolamine; PG, phosphatidylglycerol.

**Table _S32.** Similarity in polyphasic data observed between *Tepidiphilus thermophilus* JHK30$^T$ and *Tepidiphilus succinatimandens* DSM 15512$^T$. +, Positive; w+, weakly positive; −, negative; r, resistant; s, sensitive; PE, phosphatidylethanolamine; PG, phosphatidylglycerol; PC, phosphatidylcholine; DPG, diphosphatidylglycerol; AL, aminolipid; GL, glycolipid; L, unknown lipid. Data from Poddar et al. 2014

| **Characteristics** | ***Tepidiphilus thermophilus* JHK30$^T$** | ***Tepidiphilus succinatimandens* DSM 15512$^T$** |
|---|---|---|
| Biochemical tests | | |
| Urea hydrolysis | − | + |
| Reduction of nitrite | − | + |
| Activity of Enzymes (API ZYM) | | |
| Valine arylamidase | + | − |
| Cystine arylamidase | + | − |
| Trypsin | + | − |
| α-Glucosidase | + | − |
| β-Glucosidase | − | − |
| Antimicrobial susceptibility profile | | |
| Trimethoprim (5 µg) | r | s |
| Sulfamethoxazole-trimethoprim (25 µg) | s | r |
| Nalidixic acid (30 µg) | s | r |
| Growth with (as sole source of carbon) | | |
| Acetate | − | − |
| Adipate | − | − |
| Benzoate | − | − |
| l-Proline | − | − |
| Oxidation of carbon sources (Biolog) | | |
| Tween 40 | + | + |
| Tween 80 | + | + |
| i-Erythritol | − | + |
| Xylitol | − | + |
| Succinic acid monomethyl ester | + | w+ |
| *cis*-Aconitic acid | + | − |
| d-Galacturonic acid | + | − |
| d-Glucuronic acid | + | + |
| α-Ketoglutaric acid | + | − |
| α-Ketovaleric acid | − | + |
| dl-Lactic acid | + | − |
| Bromosuccinic acid | − | w+ |
| Glucuronamide | + | + |

| | | |
|---|---|---|
| d-Alanine | + | − |
| l-Alanine | − | + |
| l-Asparagine | − | + |
| l-Aspartic acid | − | w+ |
| l-Glutamic acid | + | − |
| Hydroxy-l-proline | − | w+ |
| l-Leucine | − | + |
| l-Serine | − | + |
| l-Threonine | + | w+ |
| 2,3-Butanediol | − | w+ |
| Glycerol | − | + |
| d-Glucose 6-phosphate | + | − |
| DNA G+C content (mol%) | 66.2 | 65.9 |
| Major fatty acids (>10%) | C19 : 0 cyclo ω8c, C16 : 0, C17 : 0 cyclo | C19 : 0 cyclo ω8c, C16 : 0, C17 : 0 cyclo |
| Polar lipids | PE, PG, DPG, PC, AL1, GL1–2, L1–2 | PE, PG, DPG, PC, AL1, GL1–2, L1–8 |
| Genebank accession | GCA_001418245.1 | GCA_006503695.1 |
| Genome size | 2,259,819 bp | 2,369,572 bp |

**Table S33.** Polyphasic characteristics between *Thalassospira permensis* SMB34$^T$ and *Thalassospira xiamenensis* M5$^T$ +, positive reaction or growth; –, no reaction or growth; w, weakly positive reaction; v, variable; ND, no available data. All strains are positive for assimilation D-glucose, L-arabinose, D-mannose, D-mannitol, potassium gluconate and negative for hydrolysis of gelatin, aesculin and urea, indole production, assimilation adipic and phenylacetic (API20NE test system). All strains are negative for acid production from L-rhamnose, D-melibiose and amygdalin, $H_2S$ production, lysine and ornithine decarboxylase activity (API20E test system). Similar data on acid production were obtained with conventional methods.

| **Characteristic features** | ***Thalassospira permensis* SMB34$^T$** | ***Thalassospira xiamenensis* M5$^T$** |
|---|---|---|
| Growth at: | | |
| 4 °C | − or w | + |
| 40 °C | + | + |
| 45 °C | − | + |
| NaCl range for growth (%) | 0-11.0 /0-9.0 | 0.5-10.0 |
| Optimum NaCl (%) | 2−5 | 2−4 |
| Flagellum | Monopolar single | Monopolar single |
| Nitrate to nitrite | + | + |
| Nitrate to nitrogen gas | + | + |
| Growth on carbon source: | | |
| API 20NE test system | | |
| D-Maltose | + | + |
| Capric acid | − | + |
| Citric acid | w | + |
| Malic acid | + | – |
| N-acetyl glucosamine | + | + |
| In RMM medium: | | |
| Acetate | + | + |
| D-Sorbitol | + | – |
| Aerobic acid production | | |
| D- Glucose | + | w |
| D- Mannitol | − | w |
| Inositol | − | w |
| D- Sorbitol | − | w |
| D-Sucrose | − | + |
| L-Arabinose | − | + |
| Ubiquinone's | Q-10 (Q9) | Q-10 (Q-11) |
| Accession | GCA_000300235.2 | GCA_000300235.2 |
| Genome size | 4,768,950 bp | 4,768,950 bp |
| DNA G+C contents (mol%) (genome based) | 54.7 | 54.7 |
| Major cellular fatty acids | i13: 0, i15 : 0, 16 : 0 and 16 : 1w7 (60–70 % of total), and 20 : 5w3(4–5 %). | i-15: 0, 16: 0 and 16: 1 |
| major isoprenoid quinones | Q7 (21–41 %) and Q8 (50–59 %). | |

Data from Kaster et al. 2009; Plotnikova et al. 2011.

**Table_S34.** Characteristics that distinguish *N. sicca* LMG 5290$^T$ and *N. macacae* LMG 26611$^T$. Wolfgang et al. (2013). nd, No data available.

| **Characteristic** | ***N. sicca* LMG 5290$^T$** | ***N. macacae* LMG 26611$^T$** |
|---|---|---|
| Acid production (peptone water) from: | | |
| d-Glucose | − | + |
| Maltose | + | + |
| Nitrate reduction | − | − |
| Catalase reaction | + | − |
| Acid production (API NH) from: | | |
| d-Glucose | + | − |
| Maltose | − | − |
| Sucrose | − | − |
| Enzyme activity (API NH) | | |
| Proline 4-methoxy-β-naphthylamide | + | + |
| γ-Glutamyl 4-methoxy-β-naphthylamide | − | + |
| Enzyme activity (API ZYM) | | |
| Esterase (C4) | − | + |
| Acid phosphatase | + | + |
| Major fatty acids | C16:0; C18:1ω7c; Summed feature 3 contained C16:1ω7c and/or iso-C15:0 2-OH. | C16:0; C18:1ω7c; Summed feature 3 contained C16:1ω7c and/or iso-C15:0 2-OH. |
| G+C content (mol%) | 50.9 | 50 |
| Genome size (bp) | 2,830,772 bp | 2,748,368 bp |
| GenBank Accession assembly | GCA_000174655.1 | GCA_000220865.1 |

**Table S35**. Characteristic features that distinguish *Neisseria mucosa* ATCC 19696$^T$ and *Morococcus cerebrosus* CIP 81.93T. Long et al. (1981).

| **Characteristic** | ***Neisseria mucosa*** **ATCC 19696$^T$** | ***Morococcus cerebrosus*** **CIP 81.93$^T$** |
|---|---|---|
| Cell arrangement | Irregular aggregates of 10 to 20 cells | Single, pairs, tetrads |
| Growth characteristics in microbiological medium | | |
| PYEA at 48 h | Matt colonies 1-2 mm in diameter; growth not adherent to the agar | Shiny and mucilaginous colonies 1mm diameter; growth adherent to the agar |
| Sucrose peptone agar at 48 h | Colonies 2 mm in diameter, reacting black with iodine; slime layer around cell clusters | Colonies 1 mm in diameter, black reaction with iodine extending into surrounding agar, slime layer absent |
| Growth in static PYEB | Granular | Stringy |
| Growth in shaken PYEB | Granular | Homogeneous |
| MacConkey agar | – | + |
| Acid production in Falkow amino acid basal medium | + | – |
| Reduction of litmus milk | Weak | Strong |
| G+C content (mol%) | 51.1 | 51.4 |
| Genome size (bp) | 2,688,408 bp | 2,451,909 bp |
| GenBank Accession assembly | GCA_003028315.1 | GCA_000813705.1 |

**Table S36.** Phenotypic characteristics of *Pseudoalteromonas agarivorans* KMM 255$^T$ and *P. atlantica* IAM 12376$^T$ Strains: 1, *P. agarivorans* KMM 255T, KMM 232, KMM 254 and KMM 644 (reactions in parentheses refer to strain KMM 255$^T$); 2, *P. atlantica* IAM 12376$^T$ (Akagawa-Matsushita et al., 1992). +, Positive; −, negative; v, variable between strains; w, weak; nd, not determined. Acid production was determined according to Leifson (1963). All strains were positive for the following tests: sodium-ion requirement for growth, growth at 25–28 °C, motility by a single polar flagellum, oxidase, catalase, production of lipase, caseinase, DNase, gelatin liquefaction and sensitivity to streptomycin (30 µg) and polymyxin (300 U); all strains were negative for growth at 40 °C, indole production, nitrate reduction, denitrification, arginine dihydrolase, chitin hydrolysis, l-arabinose utilization and sensitivity to lincomycin, benzylpenicillin (10 U) and O/129 (150 µg).

| **Characteristic** | ***P. agarivorans* KMM 255$^T$** | ***P. atlantica* IAM 12376$^T$** |
|---|---|---|
| Pigmentation | − | − |
| Cells with >1 polar flagellum | + | − |
| Lateral flagella | − | − |
| Growth at 4–5 °C | − | + |
| Hydrolysis of: | | |
| Starch | + | + |
| Alginate | + | + |
| Agar | + | + |
| Carrageenan | + | v |
| Acid production from: | | |
| d-Glucose | − | + |
| Maltose | v (+) | + |
| d-Mannitol | − | + |
| Oxidation of (API 20NE): | | |
| d-Glucose | w | + |
| Mannitol | + | + |
| Maltose | v (+) | + |
| Caprate | − | + |
| Adipate | − | − |
| l-Malate | − | − |
| Phenylacetate | − | v |
| Oxidation of (Biolog): | | |
| d-Fructose | − | + |
| Mannose | − | + |
| *N*-Acetylglucosamine | − | − |
| Sucrose | − | + |
| Glycerol | − | + |
| Citrate | − | + |
| Propionate | − | + |
| Butyrate | − | + |
| Glycogen | + | + |
| Sensitivity to (µg per disc):* | | |
| Ampicillin (10) | + | + |
| Gentamicin (10) | + | − |
| Kanamycin (30) | + | − |
| Carbenicillin (25) | v (−) | − |
| Oleandomycin (15) | v (+) | + |
| Tetracycline (30) | − | + |
| Neomycin (15) | v (−) | − |
| Oxacillin (20) | − | nd |
| DNA G+C content (mol%) | 40.8 | 40.8 |
| Genome size (bp) | 4,544,962 bp | 4,468,419 bp |
| GenBank Accession assembly | GCA_002310855.1 | GCA_007988745.1 |

**Table S37.** Differential phenotypic characteristics between *P. donghaensis* KCTC 22219$^T$ (Oh et al., 2011) and *P. lipolytica* JCM 15903T (Xu et al., 2010). Both strains are positive for catalase and hydrolysis of gelatin and Tween 80, and negative for lysine decarboxylase, ornithine decarboxylase and tryptophan deaminase and hydrolysis of DNA and starch. None of the strains produces $H_2S$ or indole. The following compounds are utilized as sole carbon and energy sources: N-acetyl-glucosamine, l-arabinose, maltose and mannose. The following compounds are not utilized as sole carbon and energy sources: galactose, glycerol, α-d-lactose, l-rhamnose, d-ribose, sodium malonate and sorbose. Acid is produced from maltose and mannose, but not from d-galactose, d-myo-inositol, raffinose, ribitol, α-d-lactose, sorbitol or sorbose. Both strains are susceptible to (µg per disc unless otherwise stated) amoxicillin (20), ampicillin (10), chloramphenicol (30), erythromycin (10), gentamicin (10), kanamycin (30), nalidixic acid (30), neomycin (30), penicillin (10 IU), polymyxin B (300 IU), rifampicin (5) and vancomycin (30), but resistant to bacitracin (0.04 U), clindamycin (2), nitrofurantoin (300) and nystatin (100). +, Positive; −, negative; w, weakly positive; nd, no data available.

| **Characteristic** | ***P. donghaensis* KCTC 22219$^T$** | ***P. lipolytica* JCM 15903$^T$** |
|---|---|---|
| Growth in NaCl (%, w/v): | | |
| Range | 1–13 | 0.5–15 |
| Optimum | 2–3 | 3 |
| Growth pH: | | |
| Range | 5.5–9.5 | 5.5–9.5 |
| Optimum | 6.5–7.0 | 7.0–8.0 |
| Growth temperature (°C): | | |
| Range | 4–40 | 15–37 |
| Optimum | 25–30 | 25 |
| Nitrate reduction | + | + |
| Glucose fermentation | nd | − |
| Arginine dihydrolase | − | − |
| Hydrolysis of: | | |
| Oxidase | + | + |
| Utilization of: | | |
| Sucrose | + | nd |
| Sodium citrate | + | nd |
| Acid production from: | | |
| Glucose | + | − |
| d-Fructose | nd | + |
| Mannitol | nd | − |
| l-Rhamnose | nd | − |
| d-Ribose | nd | − |
| Sucrose | nd | + |
| Trehalose | nd | − |
| Xylose | nd | − |
| Susceptibility to: | | |
| Gentamicin (10 µg) | nd | + |
| Novobiocin (30 µg) | nd | − |
| Tobramycin (10 µg) | nd | + |

| | | |
|---|---|---|
| Major fatty acids | C16 : 1 ω7c/iso-C15 : 0 2-OH, C16 : 0, C12 : 0 3-OH, SF4 (C16 : 1 ω7c and/or iso-C15 : 0 2-OH); SF7 (C18 : 1 ω7c, C18 : 1 ω9t and C18 : 1 ω12t) | C16 : 1 ω7c/iso-C15 : 0 2-OH, C16 : 0, C18 : 1 ω7c, C12 : 0 3-OH, C17 : 1 ω8c and C17 : 0. |
| DNA G+C content (mol%) | 41.6 | 41.4 |
| Genome size (bp) | 4,733,916 | 4,539,025 bp |
| GenBank Accession assembly | GCA_003515105.1 | GCA_900116435.1 |

Data from Yan et al (2016).

**Table S38.** Similarity in polyphasic data observed between *Pseudoalteromonas issachenkonii* KMM 3549$^T$ and *Pseudoalteromonas tetraodonis* IAM14160$^T$. Data from Ivanova et al. (2002). +, Positive; −, negative; ND, not detected.

| **Characteristic** | *Pseudoalteromonas issachenkonii* KMM 3549$^T$ | *Pseudoalteromonas tetraodonis* IAM14160$^T$ |
|---|---|---|
| Melanin-like pigments | - | - |
| Growth at : | | |
| 4 °C | + | + |
| 37 °C | + | - |
| Growth in NaCl at : | | |
| 8-10% | + | + |
| 12% | + | + |
| 15% | + | - |
| Production of: | | |
| Amylase | - | - |
| Alginase | + | ND |
| Agarase | - | - |
| j-Carrageenase | - | - |
| Chitinase | + | - |
| DNase | + | + |
| Utilization of: | | |
| d-Mannose | - | - |
| d-Galactose | + | + |
| d-Fructose | + | - |
| Sucrose, maltose | + | + |
| Melibiose, lactose | + | - |
| d-Gluconate | - | - |
| N-Acetylglucosamine | - | ND |
| Succinate, d-mannitol | + | - |
| Fumarate | + | ND |
| Citrate | + | + |
| m-Erythritol | ND | - |
| Glycerol | - | - |
| l-Tyrosine | ND | + |
| Xylose | - | - |
| Trehalose | - | - |
| Acetate | + | + |
| Pyruvate | + | + |
| Susceptibility to: | | |
| Kanamycin (30 ug) | - | - |
| Oleandomycin (15 ug) | + | - |

| | | |
|---|---|---|
| Ampicillin (10 ug) | + | + |
| Carbenicillin (15 ug) | + | + |
| Streptomycin (10 ug) | + | + |
| DNA G+C content (mol%) | 40.3 | 40.3 |
| Genome size (bp) | 4,131,541 bp | 4,128,400 bp |
| GenBank Accession assembly | GCA_001455325.1 | GCA_002310835.1 |

**Table S39**. Differential phenotypic characteristics of *Shewanella upenei* 20-23R$^T$ and two phylogenetically closely related *Shewanella* species Strains: 1, *S.upenei* 20-23R$^T$; 2, *S.algae* KCTC 22552$^T$. All data from this study except for motility, nitrate reduction, $H_2S$ production, urease, catalase, oxidase and DNA G+C content; these data are from Nozue et al. (1992) and Kim et al.(2007).+, positive reaction; –, negative reaction; w, weakly positive reaction. All species are Gram-negative and rod-shaped. All species are positive for motility; catalase; oxidase; $H_2S$ production, nitrate reduction; hydrolysis of casein, DNA, gelatin, tyrosine and Tweens 20, 40, 60, and 80; utilization of D-glucose, acetate, L -malate, pyruvate and succinate; activity of alkaline phosphatase, esterase (C 4), esterase lipase (C 8), leucine arylamidase (weak), Į-chymotrypsin; and susceptibility to gentamicin, kanamycin, neomycin and streptomycin. All species are negative for urease; hydrolysis of esculin, agar, starch, hypoxanthine and xanthine; utilization of L -arabinose, D-cellobiose, D-galactose, D-fructose, maltose, D-mannose, sucrose, D-trehalose, D-xylose, citrate, salicin, benzoate, formate and L-glutamate; acid production from L-arabinose, D-cellobiose, D-fructose, D-galactose, myo-inositol, lactose, maltose, D-mannitol, D-mannose, D-melezitose, melibiose, D-raffinose, L-rhamnose, D-sorbitol, sucrose, D-trehalose and D-xylose; activity of lipase (C 14), valine arylamidase, cystine arylamidase, trypsin, Į-galactosidase, ù-galactosidase, ù-glucuronidase, Į-glucosidase, ù-glucosidase, Į-mannosidase and Į-fucosidase; and susceptibility to novobiocin, cephalothin, lincomycin, penicillin G, polymyxin B and tetracycline.

| **Characteristic** | ***S. upenei* 20-23R$^T$** | ***S. algae* KCTC 22552$^T$** |
|---|---|---|
| **Acid production from** | | |
| D-Glucose | + | - |
| D-Ribose | + | - |
| **Enzyme activity (API ZYM)** | | |
| Acid phosphatase | - | + |
| Naphthol-AS-BI-phosphohydrolase | - | + |
| *N*-Acetyl-ù-glucosaminidase | - | w |
| **Susceptibility to** | | |
| Ampicillin | - | + |
| Carbenicillin | - | + |
| Oleandomycin | - | w |
| Major fatty acids | iso-$C_{15:0}$, $C_{16:0}$, $C_{16:1}$ ω7c and/or iso-$C_{15:0}$ 2-OH and $C_{17:1}$ ω8c | iso-$C_{15:0}$, $C_{16:0}$, $C_{16:1}$ ω7c and/or iso-$C_{15:0}$ 2-OH and $C_{17:1}$ ω8c |
| predominant menaquinone | MK-7 | |
| predominant ubiquinones | Q-8 and Q-7. | |
| Accession | GCA_002836995.1 | GCA_009183365.1 |
| Genome size | 4,758,780 bp | 4,866,615 bp |
| DNA G+C contents (mol%) (genome based) | 53.1 | 53 |

**Table S40.** Characteristics that differentiate *Shewanella pacifica* from phylogenetically related species: All strains are Gram-negative, motile, rod-shaped organisms that are oxidase- and catalase-positive, and can reduce nitrate to nitrite. v, Variable reaction depending on the strain; nd, data not available.

| **Characteristic** | ***Shewanella pacifica*** | ***Shewanella japonica*** |
|---|---|---|
| Growth at: | | |
| 4°C | + | – |
| 32°C | + | + |
| 0% NaCl | – | – |
| 6% NaCl | + | – |
| Haemolysis | + | + |
| Production of: | | |
| Lipase | + | + |
| Amylase | + | + |
| Gelatinase | + | + |
| Chitinase | – | – |
| Utilization of: | | |
| d-Galactose | + | – |
| dl-Lactate | – | – |
| Succinate | + | – |
| Citrate | – | – |
| GenBank Accession | GCA_003605145.1 | GCA_002075795.1 |
| Genome size | 4,831,394 bp | 4,975,677 bp |
| DNA G+C contents (mol%) (genome based) | 40.7 | 40.8 |
| Major cellular fatty acids | i13 : 0,i15 : 0, 16 : 0 and 16 : 1w7 (60–70 % of total), and 20 : 5w3(4–5 %). | i-15: 0, 16: 0 and 16: 1 |
| major isoprenoid quinones | Q7 (21–41 %) and Q8 (50–59 %). | |

**Table S41.** Characteristics of *P. elfii* SEBR 6459[T] (Ravot et al., 1995) and *P. lettingae* TMO[T] (Balk et al., 2002). Except for data on fatty acids and DNA G+C content, data are from the references listed. nd, No data available.

| **Characteristic** | ***Pseudothermotoga elfii* SEBR 6459[T]** | ***Pseudothermotoga lettingae* TMO[T]** |
|---|---|---|
| Isolation source | Oil well | Anaerobic reactor |
| Optima for growth | | |
| Temperature (°C) | 70 | 65 |
| pH | 7 | 7 |
| NaCl concentration (% w/v) | 1.2 | 1 |
| Major fatty acids* | $C_{14:0}$, $C_{16:0}$, $C_{18:0}$ | $C_{16:0}$, $C_{18:0}$ |
| Reduction of $S^0$ | − | + |
| Genome sequence accession no. | AP014507 | CP000812 |
| Genome size (Mbp) | 2.17 | 2.14 |
| DNA G+C content (mol%)† | 38.7 | 38.7 |

**Table S42.** Diagnostic and descriptive features of the two described species of *Thermotoga*. Data from Mori et al. (2014).

| **Characteristic** | ***T. petrophila*** | ***T. naphthophila*** |
|---|---|---|
| Isolation source | Oil reservoir | Oil reservoir |
| Optima for growth | | |
| Temperature (°C) (range) | 80 (47–88) | 80 (48–86) |
| pH (range) | 7 (5.2–9) | 7 (5.4–9) |
| NaCl concentration (w/v %) (range) | 1.0 (0.1–5.5) | 1.0 (0.1–6.0) |
| Major fatty acids* | $C_{16:0}$, $C_{18:0}$ | $C_{16:0}$, $C_{18:0}$, $C_{17:1}\omega 11c$, $C_{18:1}\omega 12c$ |
| Doubling time (min) | 54 | 59 |
| Motility | + | + |
| Reduction of sulfur/thiosulfate | + | + |
| Growth on sugarb | - | - |
| Mannitol | − | + |
| Xylose | – | – |
| Reduction of $S^0$ | + | + |
| Genome sequence accession no. | CP000702 | CP001839 |
| Genome size (Mb) | 1.82 | 1.81 |
| DNA G + C content (mol%)c | 46.1 | 46.1 |

37. **Kuever J, Konneke M, Galushko A, Drzyzga O**. Reclassification of *Desulfobacterium phenolicum* as *Desulfobacula phenolica* comb. nov. and description of strain SaxT

as *Desulfotignum balticum* gen. nov., sp. nov. *Int J Syst Evol Microbiol* 2001; 51:171-177.

52. **Plotnikova EG, Anan'ina LN, Krausova VI, Ariskina EV, Prisyazhnaya NV *et al.***

*Thalassospira permensis* sp. nov., a new terrestrial halotolerant bacterium isolated from a naphthalene-utilizing microbial consortium. *Mikrobiologiia* 2011; 80:691-699.

53. **Wolfgang WJ, Passaretti TV, Jose R, Cole J, Coorevits A *et al*.** *Neisseria oralis* sp. nov., isolated from healthy gingival plaque and clinical samples. *Int J Syst Evol Microbiol* 2013; 63: p.1323.
54. **Long PA, Sly LI, Pham AV, Davis GHG.** Characterization of *Morococcus cerebrosus* gen. nov., sp. nov. and comparison with *Neisseria mucosa*. *Int J Syst Bacteriol* 1981; 31: 294-301.
55. **Ivanova EP, Sawabe T, Alexeeva YV, Lysenko AM, Gorshkova NM *et al.*** *Pseudoalteromonas issachenkonii* sp. nov., a bacterium that degrades the thallus of the brown alga *Fucus evanescens. Int J Syst Evol Microbiol* 2002; 52:229-234.
56. **Mori K, Yamazoe A, Hosoyama A, Ohji S, Fujita N *et al.*** 2014. *Thermotogaprofunda* sp. nov. and *Thermotogacaldifontis* sp. nov., anaerobic thermophilic bacteria isolated from terrestrial hot springs. *Int J Syst Evol Microbiol* 64: 2128-2136.
